# In toto imaging of germ plasm dynamics reveals an essential role for early distribution of germ granules in germline development

**DOI:** 10.1101/2025.02.26.640298

**Authors:** Andreas Zaucker, Maria Papafoti, David Corcoran, DaeNia La Shawn La Rodé, Rebecca Leech, Pavle Vrliczak, Pooja Kumari, Karuna Sampath

## Abstract

A fundamental question in developmental biology is how the fertilized egg gives rise to all the different cell types of an organism. The traditional view is that the different cell types are specified either by intrinsic factors such as cell fate determinants or via intercellular signaling. In some animals a cytoplasmic determinant-like substance called ‘germplasm’ specifies the germline. In zebrafish eggs, germplasm is dispersed in form of mRNP complexes called germ granules, which are enriched at the animal pole. After fertilization the distribution of germ granules changes dramatically. The germ granules accumulate in the corners of the first two cleavage furrows of the embryo, to form four large masses that are essential for germline development. Although germ granule movement has been linked to the network dynamics of the microtubular and actin cytoskeleton, a clear mechanistic understanding of the process is currently lacking. Fundamental questions about germplasm dynamics, including “What is the main driving force?” have not been answered yet.

To address this gap, we performed careful quantitative analysis of germ granule dynamics relative to dynamic cytoskeletal reorganization in early zebrafish embryos by live-imaging. We identified stereotypic signatures of germ granule dynamics across different regions of the early embryo. Interestingly, we find that the timing of large-scale germ granule movements contrasts prevailing models for the mechanism of germ granule aggregation during cleavage divisions, and rather points to cytokinetic apparatus itself.

Using zebrafish mutants affecting the RNA-binding protein Ybx1 (Y-box binding-protein 1), we show that the timing and dynamics of germ granule accumulation in the blastodisc is a crucial factor for appropriate later aggregation into cleavage furrows and eventual distribution to PGCs. Germplasm accumulation in the cleavage furrows is reduced and ectopic aggregates form at the blastoderm margin of ybx1 mutant embryos. Our work establishes Ybx1 as a novel factor with crucial functions in germplasm distribution and suggests that additional factors drive normal germplasm dynamics.

## Introduction

How a single cell zygote gives rise to the various cell types of an organism, including the cells that give rise to the next generation, germ cells, is a fundamental question that is not fully understood. Self-organization through the establishment of signaling centers that produce growth factors and morphogens that in turn specify cell types in neighboring cells is one key strategy. A second strategy is through the action of cytoplasmic determinants, cytoplasmic factors that segregate asymmetrically during early development to control cell fate. Both strategies can specify the progenitors of the germline, primordial germ cells (PGCs), which give rise to the gametes (sperm and egg). In mice and humans, PGCs are specified by signals from neighboring cells, whereas in Drosophila, C. elegans and zebrafish, germ cells are specified by maternally provided substance called germplasm (Eno and Pelegri, 2016; Czolowska, 1969; Illmensee and Mahowald, 1974; Knaut et al., 2000; Strome and Wood, 1982),). Germplasm forms by the aggregation of a specific set of RNA-protein complexes called germ granules that are produced during oogenesis (Voronina et al., 2011). These germplasm mRNA-protein complexes (RNPs) are homotypic, i.e., the contain only one species of mRNA. RNP complexes can become part of larger assemblies such as RNP granules. Other examples of RNP complexes are various types of nuclear and cytoplasmic “bodies”(Ripin and Parker, 2023) including P-bodies, stress granules, RNA transport granules and the Balbiani body (BB) in oocytes. Studying how RNP complexes/granules assemble, function, and disassemble in living cells is therefore crucial for our understanding of gene regulation at the RNA level (Sato et al., 2022).

The study of germ granules across organisms has revealed several important insights into the biology of RNP granules. Self-sorting of RNAs into homotypic clusters within germ granules was reported in *Drosophila* oocytes (Trcek et al., 2015; Trcek et al., 2020). RNAs are translated at the germ granule surface (Chen et al., 2024; Westerich et al., 2023). Gradients of molecular activities were found to bias germ granule condensation to the posterior half of polarized C elegans zygotes (Wang et al., 2014; Wu et al., 2019). In early zebrafish oocytes, the centrosome nucleates the formation of the Balbiani body (Elkouby et al., 2016). The BB in Xenopus and zebrafish has an amyloid-like character and form by phase-separation of proteins that carry Prion-like domains, Bucky ball and Xvelo respectively (Boke et al., 2016; Deis et al., 2022). Although new insights have been obtained from these studies, fundamental questions about the distribution and dynamics of germ granules in vertebrate embryos remained unanswered.

Germ granules are enriched at the animal pole of the zebrafish egg, the side where the cytoplasm will accumulate after egg activation and fertilization to form the blastodisc where the embryo will develop. The initially dispersed germ granules accumulate in the corners of the cleavage furrows during early cleavage divisions and form large masses of germplasm, also called furrow aggregates. The masses become progressively smaller with each cleavage division and only aggregates found in the cleavage furrows from the first two divisions are thought to have the critical mass to specify primordial germ cells (PGCs) later during embryonic development (Moravec and Pelegri, 2020). Four large masses of germplasm form and are later taken up by four cells, which eventually will give rise to the germline (Yoon et al., 1997). The formation of the four clusters masses/aggregates is an essential step during germline development in zebrafish, and their experimental removal in embryos leads to sterility in adults (Hashimoto et al., 2004).

Much of what we currently know about the mechanism of germplasm aggregation during early cleavage divisions has been inferred from the analysis of zebrafish mutant phenotypes and the effects of interference with the cytoskeleton by *in situ* staining methods (Eno and Pelegri, 2013; Eno and Pelegri, 2018). Hence, our current knowledge about this highly dynamic process is largely based on snapshots of what is likely a highly dynamic process.

While it seems clear that the embryo uses the cell division machinery to collect germ granules in the corners of the cleavage furrows, the exact molecular mechanism is still elusive. Key questions like “what is the main driver of germ granule movements” are currently still without a definitive answer. Quantitative analysis of germ granule dynamics during early cleavage divisions is crucial to advance our understanding of what drives their movement and aggregation.

To quantitatively characterize germ granule behavior prior to and during the first cell divisions, we performed time-lapse imaging of the germplasm/germ granule reporter Buc-EGFP (Riemer et al., 2015; Zaucker et al., 2021). We identified signature behaviors of various subpopulations of germ granules based on their initial localization within the embryo. Interestingly, we found that current models for the mechanism of germ granule dynamics cannot be reconciled with some of our findings. Our work links bulk germ granule dynamics to the formation and ingression of the cleavage furrow, and the cytokinetic apparatus. Mutation of a gene that encodes for a protein that has been implicated in cytokinesis, Y-box binding protein 1 (Ybx1), leads to sex ratios biased towards males in adults, and to reduced numbers of PGCs in maternal *ybx1* mutant embryos. Our investigation into the causes of the germline phenotype in *ybx1* mutants revealed that the furrow aggregates are smaller in *ybx1* mutants than in controls. Surprisingly, this is a consequence of altered germ granule dynamics and distribution in the mutants, rather than reduced germplasm in the mutants.

Based on our findings, we posit that a delay of germ granule movement towards the animal pole in *ybx1* mutant embryos results in smaller furrow aggregates and the formation of ectopic aggregates. This results in reduced numbers of PGCs in mutant embryos and likely underlies the male bias in ybx1 mutants. Our work identifies Ybx1 as a novel factor that affects germ plasm distribution and dynamics in early zebrafish embryos and provides evidence for a role for early germplasm distribution in regulation of later events in germline development.

## Results

### Germ granule movement is stereotypical for distinct regions of the blastodisc

To study germ granule dynamics during early embryonic cell divisions, we recorded the movement of germ granules by time-lapse imaging of embryos of the Tg(buc:buc-egfp) germplasm/granule reporter line (Figure1A). We combined *in toto* imaging of the entire blastoderm at low magnification on a spinning disc confocal microscope (SDM; 717×717×150 μm volumes at 20-30 s intervals), with imaging of a small area at high spatiotemporal resolution (106×106×53 μm volumes, 1 s intervals) on a lattice lightsheet microscope (LLSM, Figure 1B).

**Figure 1.**
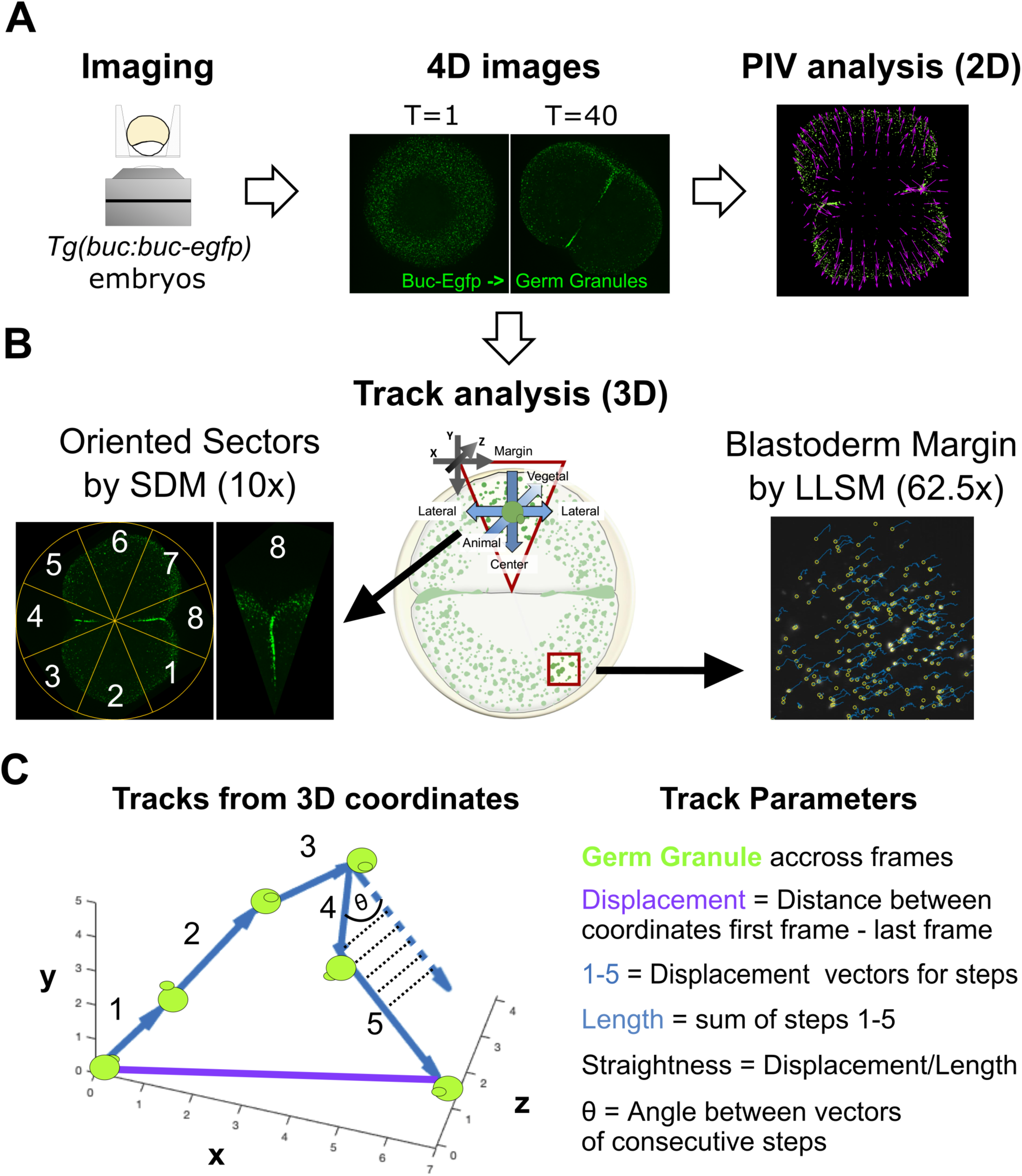
Outline of quantitative characterization of germ granule dynamics. **A)** Workflow to analyze germ granule dynamics by particle imaging velocimetry (PIV). Mounting tools were used for time-lapse imaging of Buc-Egfp labelled germ granules during early cell divisions and PIV analysis to construct vector maps of the flow of the germ granules. **B)** Imaging of germ granule movement across scales by 3D tracking of individual granules. Left, principle of standardized parameter measurement of germ granule dynamics from animal pole views of *in toto* imaging by spinning disc microscopy. Segmentation of imaged embryos with the 1^st^ cleavage furrow oriented horizontally generates numbered, equal-sized sectors, covering comparable regions across different embryos. Reorientation of all numbered sectors with the pointed end (red outlined triangle on the schematic at the center) aligning the coordinate system of sector volume (top left corner) with defined axes within the embryo (center of triangle). Right, high spatiotemporal imaging of a small region near the blastoderm margin (red square on the schematic at the center) on a lattice light sheet microscope to estimate parameters at high precision. **C)** Schematic outlining how the 3D coordinates of germ granules derived from 3D tracking were used to calculate germ granule kinetics (right).

In the *in toto* imaging movies we observed only modest movements of germ granules in the minutes prior to the first cell division (cleavage), with no obvious regional differences. Following the appearance of the first cleavage furrow, germ granules move into the furrow, but only from areas adjacent to the furrow (Movies1,2).

To investigate the differences in germ granule behavior across the embryo, we compared parameters of germ granule dynamics between subpopulations of germ granules from different regions (sectors) within the imaged embryos (Figure1B). Parameters were measured by particle image velocimetry (PIV) and 3D track analysis (Figure1A and C). For those experiments we used Tg(buc:buc-egfp); *Tg(actb2:mCherry-Hsa.UTRN)* double-transgenic embryos. The Tg(actb2:mCherry-Hsa.UTRN) transgene encodes for a f-actin reporter that facilitated detection of the cleavage furrows (Figure 2, Movie2).

**Figure 2.**
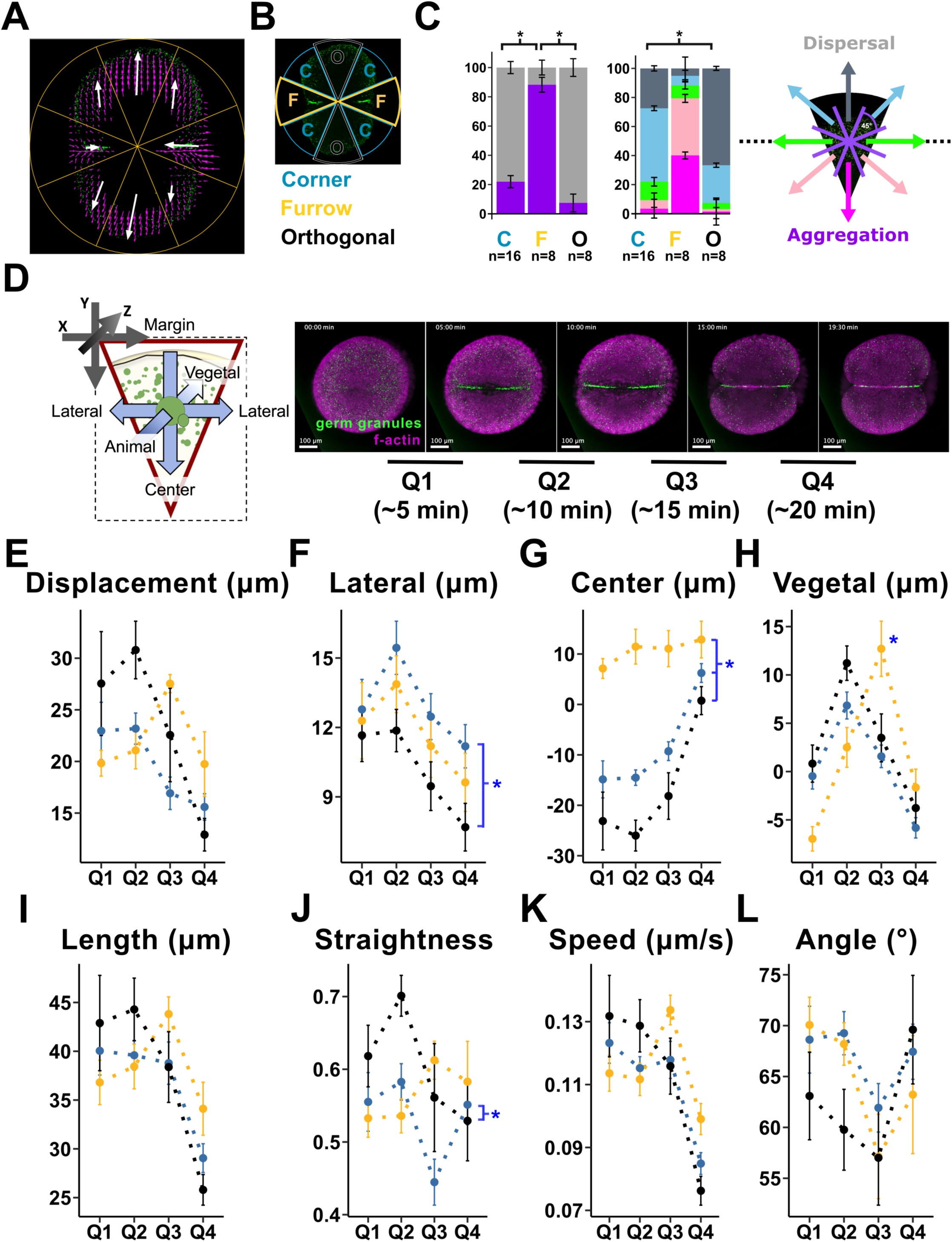
Germ granule movement is highly stereotypic for different positions within the blastodisc. **A)** 2d vector map for average movement of Buc-Egfp positive germ granules (green) during the 1^st^ cleavage division from the PIV analysis of a time-lapse recording in Tg(*buc:buc-egfp*) transgenic embryo by spinning disc microscopy. Outlines of sectors are overlayed in orange; purple arrows, individual vectors; white arrows, summary vectors for the sectors (average). **B)** The 1^st^ cleavage furrow generates bilateral symmetry of the embryos in animal pole views. Standardized segmentation of the embryo with the (future) cleavage furrow in horizontal orientation produces groups of opposing sectors that cover comparable regions of the embryo: Furrow (F, golden), Orthogonal (O, black), and Corner (C, steel blue) between F and O. This color scheme is used throughout the figure and manuscript. **C)** Quantitation of movement directionality of germ granules in different sector groups by calculation of the proportions of vectors falling into different categories for directionality. Categories and corresponding range of vector angles are indicated by color code (legend on right): margin (slate-grey), diagonal up (sky blue), lateral (green), diagonal down (pink), center (magenta). Movement towards the margin (margin, diagonal up) was combined into a category called “Dispersal” (light grey), whereas movement towards the center (diagonal down, center) or into the furrow (lateral) was combined into a category called “Aggregation” (purple). The stacked bar chart on the left compares ‘dispersal’ vs ‘aggregation’ movements between different sector groups. Stacked bar chart (right) compares vectors for all directionality categories between sector groups. **D)** Left, relationship between the coordinate system of the reoriented sectors during 3D tracking of germ granules and the movement of germ granules along defined axes within the embryo calculated from the tracks. Top left, origin of the 3D cartesian coordinate system of the reoriented sector movies (4D images). The dimensions of the 4D image of the cropped and reoriented sector are outlined with dashed lines and the cropped sector is depicted as a red triangle. At the center of the sector is a large germ granule (green) together with the coordinate system for its movement along defined axes within the embryo at any given time interval. On the right, temporal segmentation to obtain temporal profiles for the parameters. The time-lapse movie of the first division was divided into four segments, Q1-Q4. Stills demarcating the different segments are arranged sequentially from left to right. **E-L)** Parameter profiles describing germ granule movement across the time periods in D.

We identified three different groups of sectors based on the PIV analysis of the movies (Figure 2A,B). In non-furrow sectors (groups C and O), the summary vectors (white) point towards the margin, whereas in furrow sectors (group F) they point towards the center of the embryo. The two largest summary vectors are the ones from the opposing sector pair (group O) that is perpendicular to the furrow-sector pair (group F). This differentiates the orthogonal sectors (group O) from the corner sectors (group C) adjacent to the furrow-sectors (group F). Corner sector vectors are smaller and less aligned towards the margin, presumably due to the influence of the cleavage furrow.

Quantitative analysis of the directionality of individual velocity vectors (pink) for each sector of the movies corroborated this finding. On the vector maps, arrays of opposing vectors flanking a string of vectors pointing towards the center of the cell can be seen in the furrow sectors. The furrow sectors have the highest percentage of sectors oriented diagonally down (39%) and towards the center (40%), together with some vectors pointing laterally (9%). This reflects the movement of germ granules into the cleavage furrow seen in the movies.

Therefore, we reasoned that these three vector directions reflect movements that contribute to the aggregation of germ granules. In contrast, movements towards the margin, either straight or diagonally, disperse germ granules and render them unavailable for aggregation (Moravec and Pelegri, 2020). The data for these two categories, “aggregation” versus “dispersal”, are clearly distinct between furrow and non-furrow sectors. Quantitative analysis shows differential “aggregation” in furrow sectors (88%), corner sectors (22%) and orthogonal sectors (7%).

The proportions of vectors pointing straight towards the margin (margin) versus diagonally up towards the margin (diagonal up) distinguish between corner and orthogonal sectors. Most vectors in orthogonal sectors (67%) point straight towards the margin. The vectors in the corner sectors also point towards the margin but the majority are in a diagonal orientation (51%), with only 27% of vectors pointing straight towards the margin (Figure2C).

PIV analysis revealed three stereotypic patterns of germplasm dynamics across the sectors: (1) into the furrow and towards the center of the embryo (furrow sectors, F), (2) highly coordinated movement towards the blastoderm margin (orthogonal sectors, O), and (3) less-coordinated movement towards the margin (corner sectors, C). However, PIV analysis does not provide movement dynamics of individual granules as it treats germplasm as a fluid and generates vector maps for flow.

### Differential movement towards vegetal in different regions of the blastodisc

To determine how individual granules move, we performed 3D track analysis. We combined the spatial segmentation of the embryo into sectors with temporal segmentation of ∼20 minutes (min) movies into four equal-sized segments (Q1-4 in Figure 2D) to obtain temporal profiles for the parameters of germ granule movement in the sectors in 3D: displacement (total, center, lateral, vegetal), length, straightness, speed, and angle between consecutive 3D vectors. In contrast to summary statistics obtained from analysis of the whole movie, these profiles contain additional information about dynamic changes in the movement of germ granules during the first cleavage division. Figure 2E-L summarizes the profiles.

Our analysis did not reveal a significant difference in the speed of germ granules between sector groups during the first cleavage division (Figure 2K). Accordingly, there was also no significant difference in the length of the tracks (Figure 2I). This was also the case for the 3D angles (Figure 2L).

Analysis of the individual components of the total displacement vectors (Figure 2E-H) confirmed the stereotypic patterns we saw in the vector maps of PIV analysis. The granules move towards the center in furrow sectors (positive values), and towards the margin (negative values) in non-furrow sectors (Figure 2G). All sector groups are significantly different from the other groups for this parameter (C vs O p-value = 0.0005, C vs F p-value = 0, C vs O p-value = 0). The profiles show that the movement towards the margin mainly happens during the first three segments Q1-3 in the non-furrow sectors, while there is a continuous displacement towards the center throughout in the furrow-sectors.

The 3D tracking data also validated the finding that the germ granules follow less straight trajectories towards the margin in corner sectors. Several parameters related to this observation were significantly different between corner and orthogonal sectors (Lateral p-value = 0.004, Straightness p-value = 0.04).

The movement along the animal-vegetal axis sharply peaked during one segment in all sector groups (Figure 2H). However, the peak shifts from Q2 (corner = 6.8 μm, furrow = 2.5 μm, orthogonal = 11.2 μm) in non-furrow sectors to Q3 in furrow-sectors (corner = 1.6 μm, furrow = 12.7 μm, orthogonal = 3.5 μm). Similar results were found with single transgenic embryos harbouring the Tg(buc:buc-egfp) transgene (Figure S1).

Together, the analysis of the movement of germ granules in the different sector groups validated the patterns identified by PIV analysis. However, the inclusion of the third dimension (animal-vegetal), revealed an additional difference between furrow and non-furrow sectors. This suggests that the furrow itself might be a key factor influencing germ granule behaviour.

### Wave of displacement during 1^st^ division

The 3D track analysis showed that germ granule movement towards vegetal differs in regions outside the furrow versus at the furrow. To investigate this in-depth, we analysed germ granule dynamics with respect to furrows (animal pole, Figure3A and Movie3; lateral views, Figure3B and Movie4).

In color-coded movies, germ granules in the cleavage furrow start moving through colors of increasing z-depth at segment Q3 (Figure3C, Movie3). The lateral view movies show that germ granules outside of furrow region get displaced towards the blastoderm margin in form of a wave (outward wave) emanating from the cortical center of the blastodisc (Figure 3D). In contrast, vegetal movement of germ granules near the furrow mirrors furrow ingression and is delayed relative to the outward wave. In addition, some germ granules are not incorporated into the outward wave and largely remain stationary.

**Figure 3.**
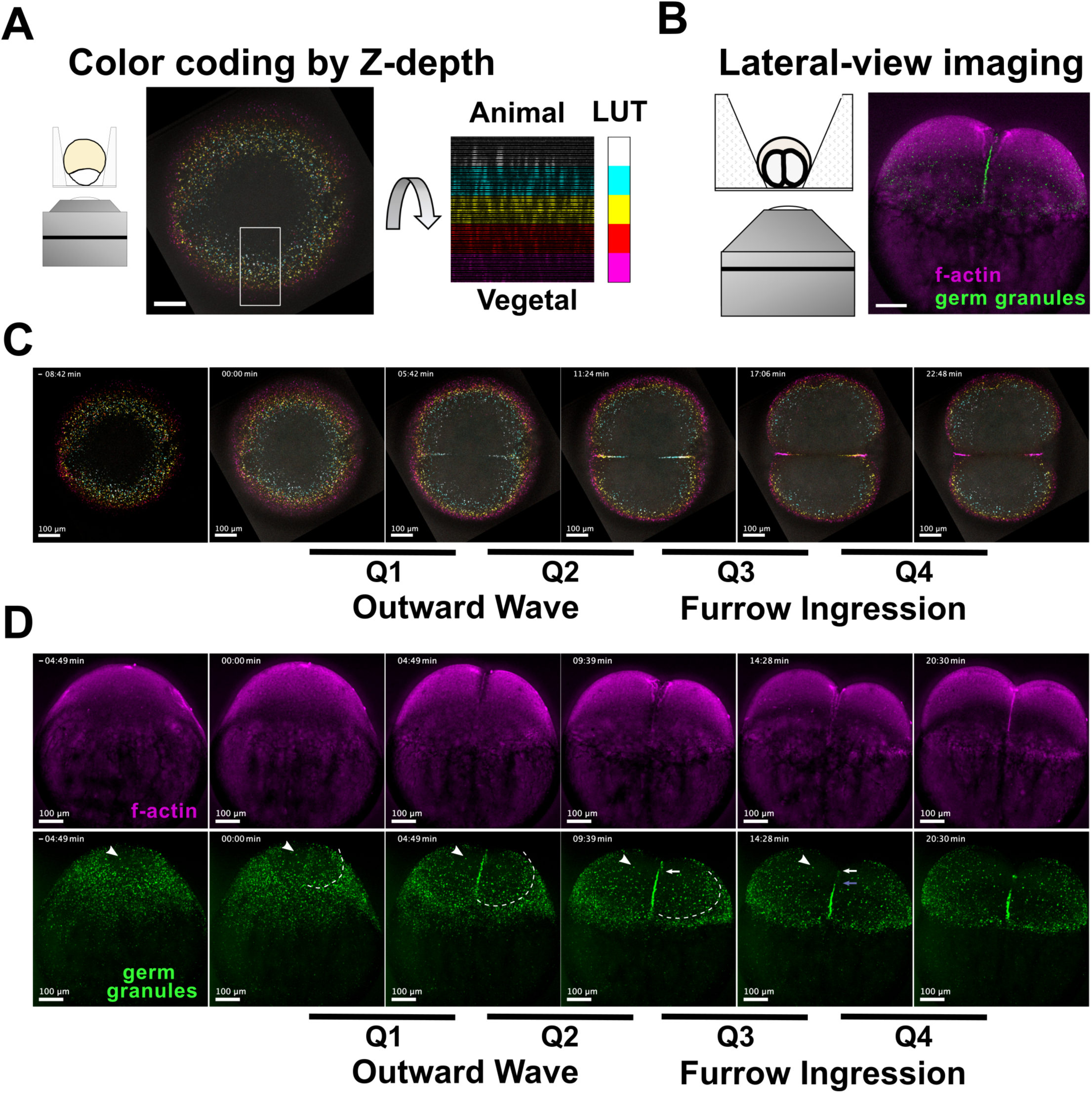
Wave of displacement during 1^st^ division, with some granules remaining stationary. **A)** Left, schematic of mounting strategy for animal pole views. Right, color-coding to visualize positional information of germ granules along z in maximum intensity projections (MIPs). **B)** Left, schematic of the mounting strategy for lateral views. Right, still from a time-lapse movie of f-actin (Utrn-mCherry, magenta) and germplasm (Buc-EGFP, green) during the 1^st^ embryonic cell division. **C)** Series of stills (left to right) from time-lapse imaging of Buc-Egfp germplasm during the 1^st^ cleavage. The first still is from a timepoint just before the cleavage furrow appears. Other stills demarcate four equal-timed segments of the 1^st^ division (Q1-Q4). Z-depth of the Buc-Egfp signal was color-coded with the same LUT as in A. **D)** Stills from time lapse imaging during the first division (lateral views). The sequence of stills is aligned to that in panel C. The top row shows the signal for Utrn-mCherrry (f-actin, magenta), and the bottom row shows the signal for Buc-EGFP (germplasm, green). White arrowheads point to a granule which largely remained stationary during imaging. The front of a wave of germ granule displacement towards the margin is indicated by a dashed line. The white arrow points to the top position regarding the animal-vegetal axis of the “near” furrow aggregate at the beginning of segment Q3, while the blue arrow points to its position at the end of Q3. All scale bars = 100 μm.

### Large-scale movement of germ granules only commences with cleavage furrow formation

To pinpoint the exact time-point when the outward wave is initiated, we imaged the germplasm reporter Tg(buc:buc-egfp) together with the f-actin reporter Tg*(actb2:mCherry-Hsa.UTRN)* in animal pole and in lateral views (Figure 4, Movies5,6). Before furrow formation, an actin polymerization wave runs over the cortex without substantial changes in germ granule distribution (Figure 4 A,B,D, E). With the appearance of the furrow (contractile band), dramatic changes occur in the distribution of germ granules in the form of an outward wave (Figure 4C and F). These results suggest that large scale movement of germ granules is only begins after cytokinetic cleavage furrow initiation.

**Figure 4.**
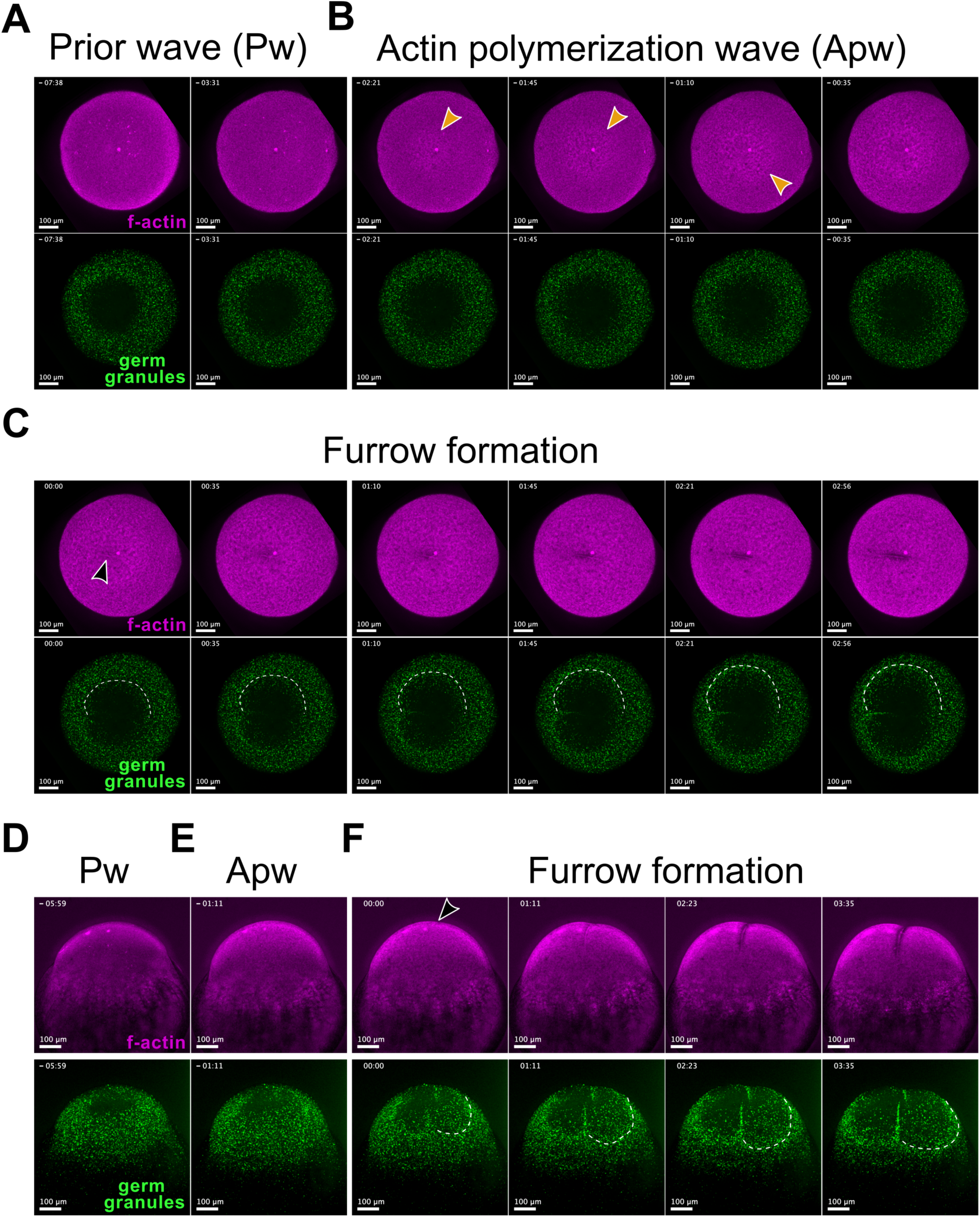
Germ granule movements increase with appearance of the 1^st^ cleavage furrow -. Stills from time lapse spinning disk microscopy imaging of Tg(buc:buc-egfp); Tg(actb2:utrn-mcherry) double transgenic embryos showing Buc-EGFP labelled germ granules(green), Utrn-mCherry labelled f-actin (magenta). Panels A-C, animal pole views of an embryo, panels D-F show another embryo in lateral views. **A,D)** Prior to furrow formation, the f-actin signal is relatively weak and only modest movement of germ granules is observed. **B,E)** An actin polymerization wave over the embryo has little impact on germ granule distribution. **C,F)** Actin is polymerized. A wave of germ granule displacement towards the margin is seen with formation of the cytokinetic band (black arrowhead). Furrow formation and wave progression in parallel throughout consecutive frames.

### Maternal ybx1 mutants have a germline defect

We then deployed our method to detect changes in germ granule dynamics after perturbations (chemically, genetic or physical) of cellular factors and networks, to identify potential novel factors that govern germ granule dynamics.

Ybx binding-protein 1 is a nucleic acid binding protein with roles in regulation of gene expression and the cytoskeleton (Chernov et al., 2008; Kawaguchi et al., 2015; Mehta et al., 2020; Ruzanov et al., 1999; Sun et al., 2018; Uchiumi et al., 2006). Hence, we investigated germline development in a temperature-sensitive ybx1 mutant allele, *ybx1^sa42^* (Kumari et al., 2013). Henceforth, we refer to homozygous mutants for the *sa42* allele as *ybx1* mutants.

Families of fish raised from clutches laid by homozygous *ybx1* mutant mothers displayed a male sex bias (Figure 5A, Figure S2B). This was observed in embryos derived from crosses of mutant parents, i.e. maternal zygotic *ybx1* mutants (MZ*ybx1*, 59% males) which lack maternal and zygotic Ybx1 protein. Importantly, ybx1 mutants from crosses of *ybx1* mutant females to wild type (wt) males, that only lack maternal Ybx1 protein, i.e. M*ybx1* mutants, had a similar male bias (63% males). Families of control fish did not show a male bias.

**Figure 5.**
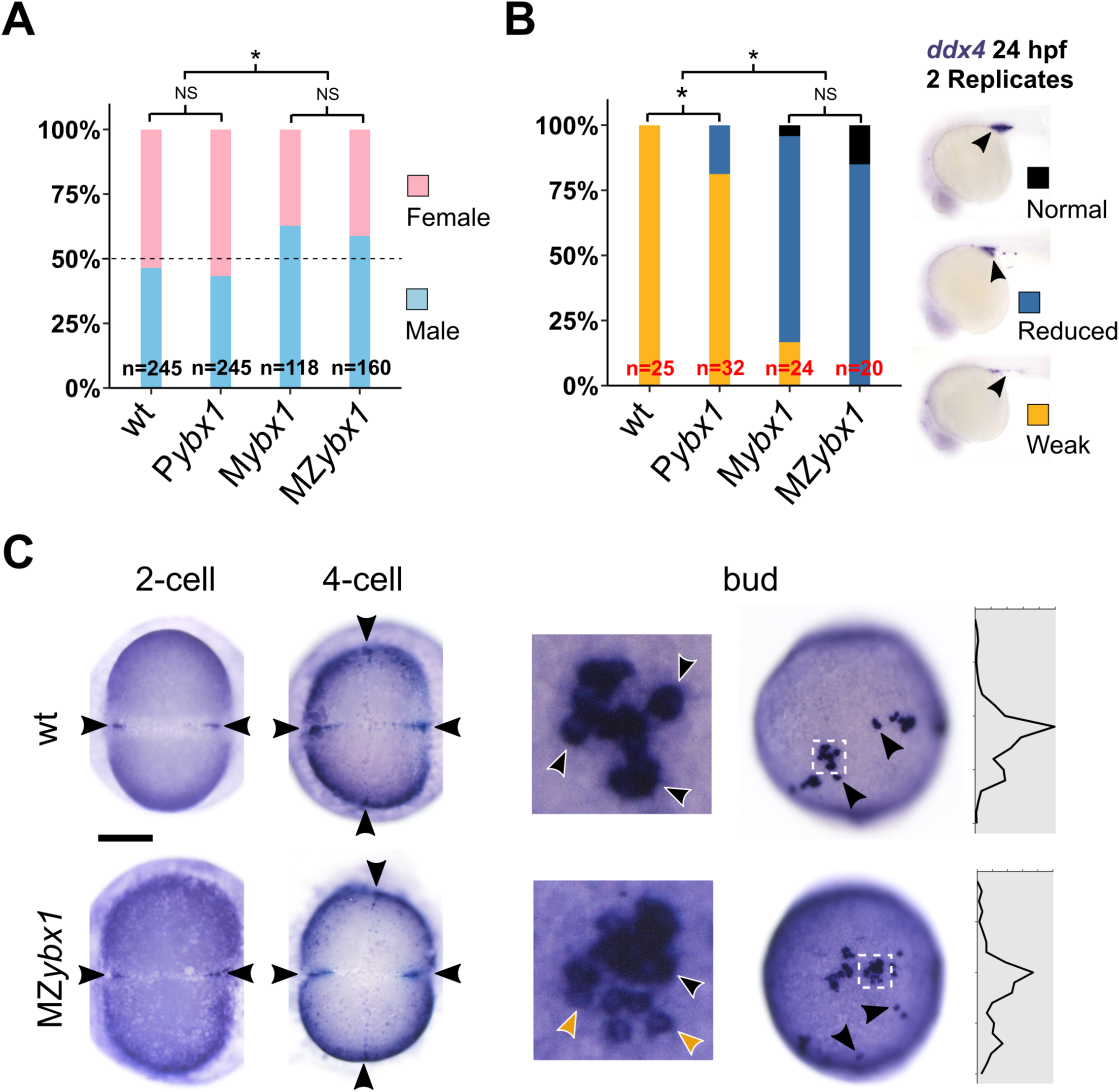
Maternal ybx1 mutants have germline defects. **A)** Male versus female sex ratios in families of adult fish of various genotypes. **B)** Stacked bar chart shows % of 24 hpf embryos per genotype and category of *ddx4* expression. Representative images for the various categories and corresponding color code are shown on the left. Arrowheads indicate PGCs. **C)** Stage series shows WISH to detect *ddx4* in MZ*ybx1* mutants versus wild type (wt) siblings during early development. Animal pole views shown, except for bud stage (lateral view with animal to the top). Histograms (left) show PGC distribution along the animal vegetal axis. Arrowheads indicate position of germplasm aggregates in 2-cell and 4-cell, and of clusters of PGCs at bud stage; area indicated by dashed box shows individual PGCs. Yellow arrowheads indicate weakly stained PGCs in MZ*ybx1* mutants. Black, control/normal and dark gold for mutant/abnormal.

Male sex bias can arise from reduced numbers of PGCs or reduction in germplasm (Tzung et al., 2015). To investigate if the male bias in ybx1 mutants arises from reduced germplasm and/or PGC numbers, we performed whole-mount RNA in situ hybridization to detect vasa/*ddx4* expression in PGCs of 24 hpf embryos. Expression of *vasa/ddx4* is reduced in MZ*ybx1* and M*ybx1* mutants compared to wt and P*ybx1* controls (Figure 5B) raised at 22 or 28°C (Figure S2C-E). The number of cells in the gonadal region carrying the Buc-Egfp-labelled germplasm is reduced in Tg(*buc:buc-egfp*) transgenic MZ*ybx1* mutants in comparison to controls derived from crosses of Tg(*buc:buc-egfp*); ybx1^sa42/+^ females with wt males (Figure S2F,G).

To determine when germplasm is lost in ybx1 mutants, we examined *vasa/ddx4* expression in a stage series of embryos. MZ*ybx1* mutant embryos initially have germplasm that accumulates in the corners of the first cleavage furrows in a pattern similar to wt controls (Figure 5C). At bud stage, a slight difference in the distribution of PGCs between MZybx1 mutants and controls can be discerned. While PGCs migrate in tight columns of cells in wt controls, the mutant PGCs seem slightly more scattered. Additionally, some PGCs in *ybx1* mutants show weaker staining than wt controls. In summary, maternal ybx1 mutant embryos show a bias towards males, and the number of PGCs colonizing the gonadal region is slightly reduced.

### Germplasm distribution is altered in MZ*ybx1* mutants by early cleavage stages

Although we observed similar overall expression of germplasm RNAs *ddx4* and *dnd1* in 4-cell stage MZybx1 and wt embryos, the WISH staining intensity seemed reduced, especially in the cleavage furrows (Figure 6A). This suggests maternal *ybx1* mutant eggs might contain less germplasm. To test this, we performed comparative gene expression analysis on 4-cell stage MZ*ybx1* and P*ybx1* control embryos. Transcripts for *ddx4* and *dnd1* did not appear reduced in Mybx1 mutants by qPCR. Analysis of RNAseq experiments validated the qPCR data and did not show a change in germline transcript levels in mutant embryos (Figure 6B). Together, the gene expression analysis in 4-cell stage embryos suggested altered distribution of germplasm without changes in RNA expression levels.

**Figure 6.**
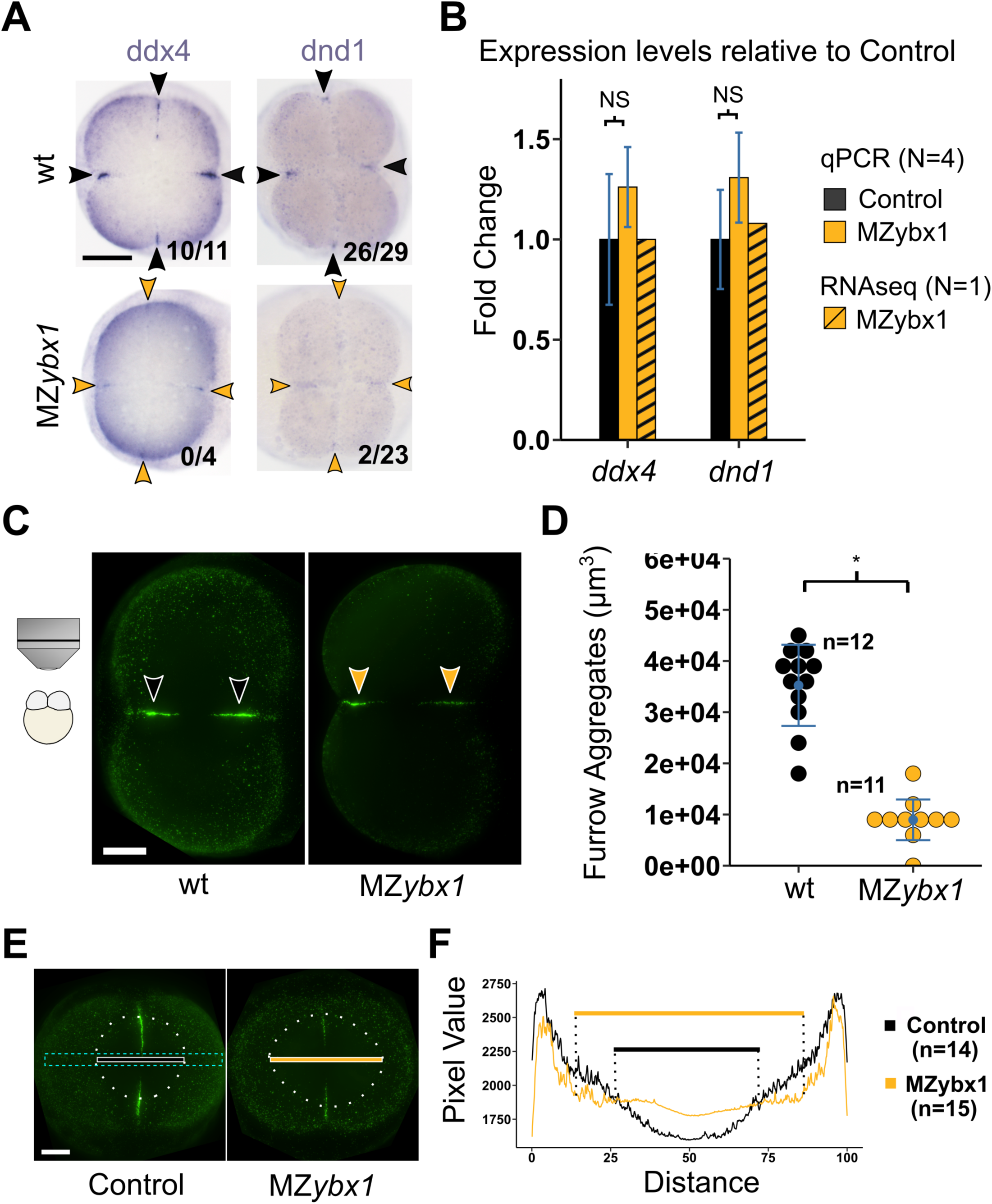
Germplasm distribution is altered in 2-4 cell stage MZ*ybx1* mutant embryos. **A)** Representative images of WISH to detect *ddx4* and *dnd1* in 4-cell embryos. Black arrowheads point to germplasm masses in the corners of the cleavage furrows of wt embryos, dark gold arrow heads indicate smaller masses seen in *ybx1* mutants. Scale bar = 200 μm. **B)** Quantitation of *ddx4* and *dnd1* expression in *ybx1* mutants versus controls. Expression levels were normalized to 18S RNA;error bars indicate SEM. Dashed bars indicate fold change for the same genes from RNAseq. **C)** Representative 2-cell stage embryos showing Buc-EGFP labelled germplasm in MZ*ybx1* mutants versus wt controls; Scale bar = 100 μm. **D)** Dot plot of the Egfp+ volumes of the cleavage furrow aggregates in embryos from panel C, obtained by 3D segmentation after deconvolution. **E)** Measurement of germplasm distribution across the blastoderm from animal pole views of 2-cell embryos by widefield microscopy. The embryos are oriented with the cleavage planes vertically. Germ granule-free zones in mutants and controls are outlined by dashed ellipsoids (white), with the diameter represented by black (control) and dark gold (mutant) lines. 100 pixel-wide line scans were measured across the area outlined by a dashed turquoise rectangle. Scale bar, 100 μm. **F)** Summary line graphs showing the average EGFP-signals for mutants and controls at comparable positions across the blastoderm in 0.1% bins of the total lengths of the line scans.

Analysis of a germplasm protein, Bucky ball, in Tg(*buc:buc-egfp*) mutant transgenics showed that the volume of the EGFP+ germplasm aggregates in the corners of the first cleavage furrow was reduced in MZybx1 embryos compared to controls (wt = 3.5e+4 μm^3^, MZ*ybx1* = 0.9e+4 μm^3^). This is consistent with our germplasm WISH findings (Figure 6C,D, Figure S3A-D).

To test if the distribution of germ granules is altered, we measured Buc-eGFP signal intensity by line scans as a proxy for germ granules, across mutant and control embryos. The germplasm free zone at the center of the blastodisc in animal pole views seems enlarged in MZ*ybx1* embryos (Figure 6E). The histogram for the average EGFP signal at normalized positions on the line scans shows that germ granules are differentially distributed in mutants (Figure 6F).

### Germplasm dynamics are altered in MZ*ybx1* mutants

We hypothesized that changes in the dynamic behavior of germ granules might underly the altered distribution of germ granules in *ybx1* mutants compared to wt embryos. To test this, we used our pipeline for comparative analysis of germ granule movement dynamics between wt and MZ*ybx1* mutant embryos (Figure 7).

**Figure 7.**
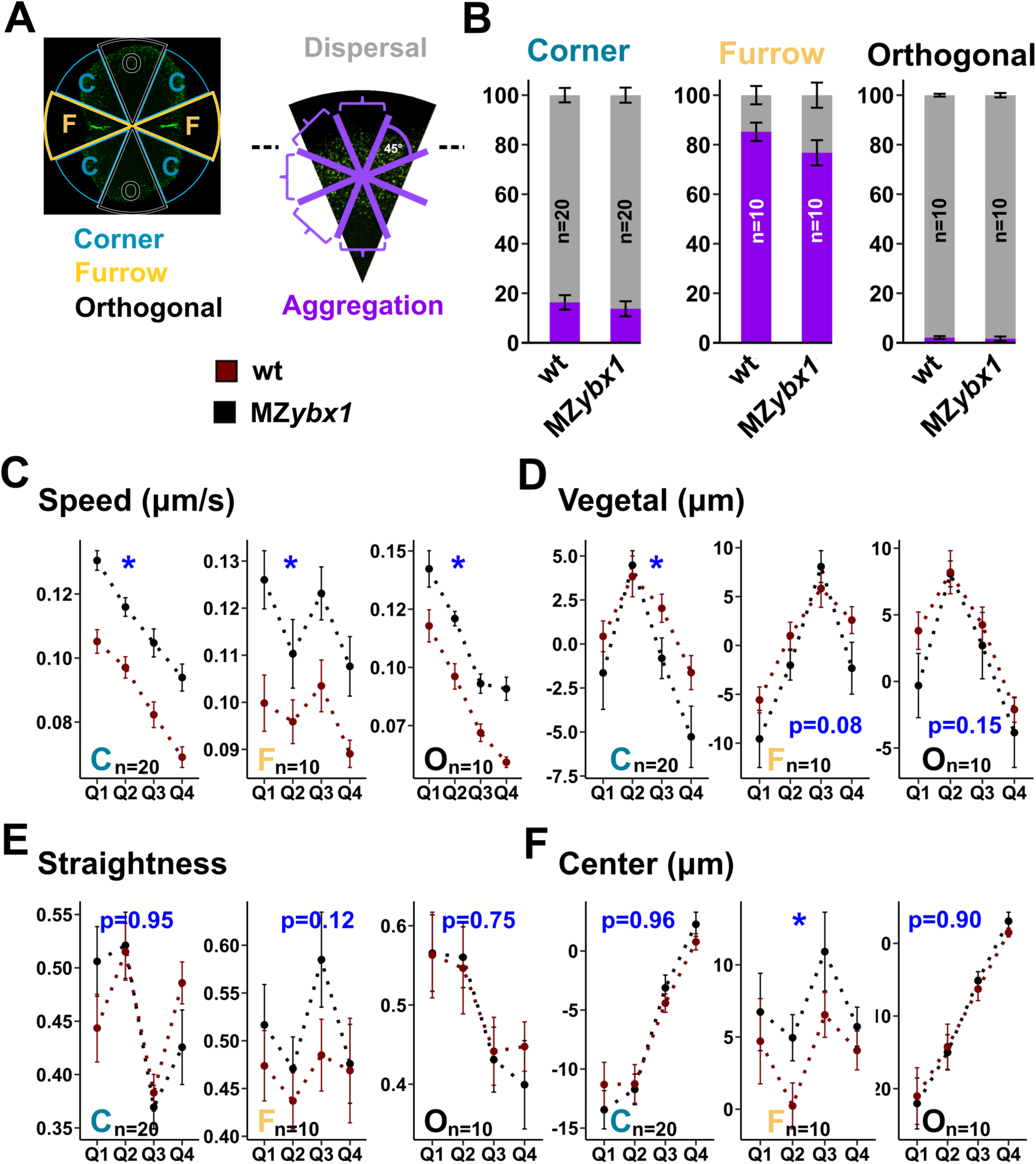
Germplasm dynamics are changed in MZ*ybx1* mutants. **A)** Legend for panels B-F. Schematic on left shows the different sector groups, with names and color code listed below; Right, a sector in standard orientation illustrating the range of vector directions in “Dispersal” and “Aggregation” categories. In line graphs C-F, black data points indicate wt and red data points indicate MZ*ybx1* mutants. **B)** Stacked bar charts comparing % vectors with Dispersal vs Aggregation directionality across different sector groups in wt versus mutants. **C-F)** Line graphs comparing the temporal parameter profiles of germ granule dynamics across different sector groups in wt versus MZ*ybx1* mutants. P-values from two-way ANOVA are given in blue font (asterisks indicate p < 0.05).

Aggregation-related vector directionality was slightly reduced in the furrow and corner sectors of *ybx1* mutants. In furrow sectors, 85% of vectors (n=10) in wt embryos fall into the “aggregation” category, versus 77% in mutants (n=10). In corner sectors 16% aggregation was observed for wt (n=20) and 14% in MZ*ybx1* mutants (n=20). However, the observed differences were not statistically significant.

PIV analysis indicates changes in germ granule movements in *ybx1* mutants, specifically in the furrow region. This was corroborated by the results of the 3D track analysis (Figure 7C-F). The speed (Figure 7C, Figure S4) and length of the tracks was reduced in the mutants across all sector groups. Additional significantly different parameters were only observed in the furrow region (furrow and furrow-adjacent corner sectors): Displacement (furrow p-value = 0.018, corner p-value = 0.007), Vegetal (corner p-value = 0.03) and Center (furrow p-value = 0.03). Interestingly, for several other parameters the p-values were very low in furrow sectors (Vegetal p-value = 0.08 and Straightness p-value = 0.12).

Taken together, our germ granule analysis method detected differences in germ granule dynamics, particularly regarding the speed of movement, between ybx1 mutants and wt controls. Furthermore, the differences were more pronounced in the furrow region.

### Germ granules mis-aggregate at the blastoderm margin in MZ*ybx1* mutants

To investigate the cause of altered germplasm distribution in mutant embryos, we recorded movies of the movement of germ granules prior to and during the first cleavage divisions by confocal spinning disc microscopy. In animal pole and lateral views (Figure 8A-D, Movies7-10), we found that blastodisc elevation is reduced in the mutants. We also observed substantial germplasm aggregation at the blastoderm margin rather than only in the cleavage furrows in the mutants (Figure 8B,D).

**Figure 8.**
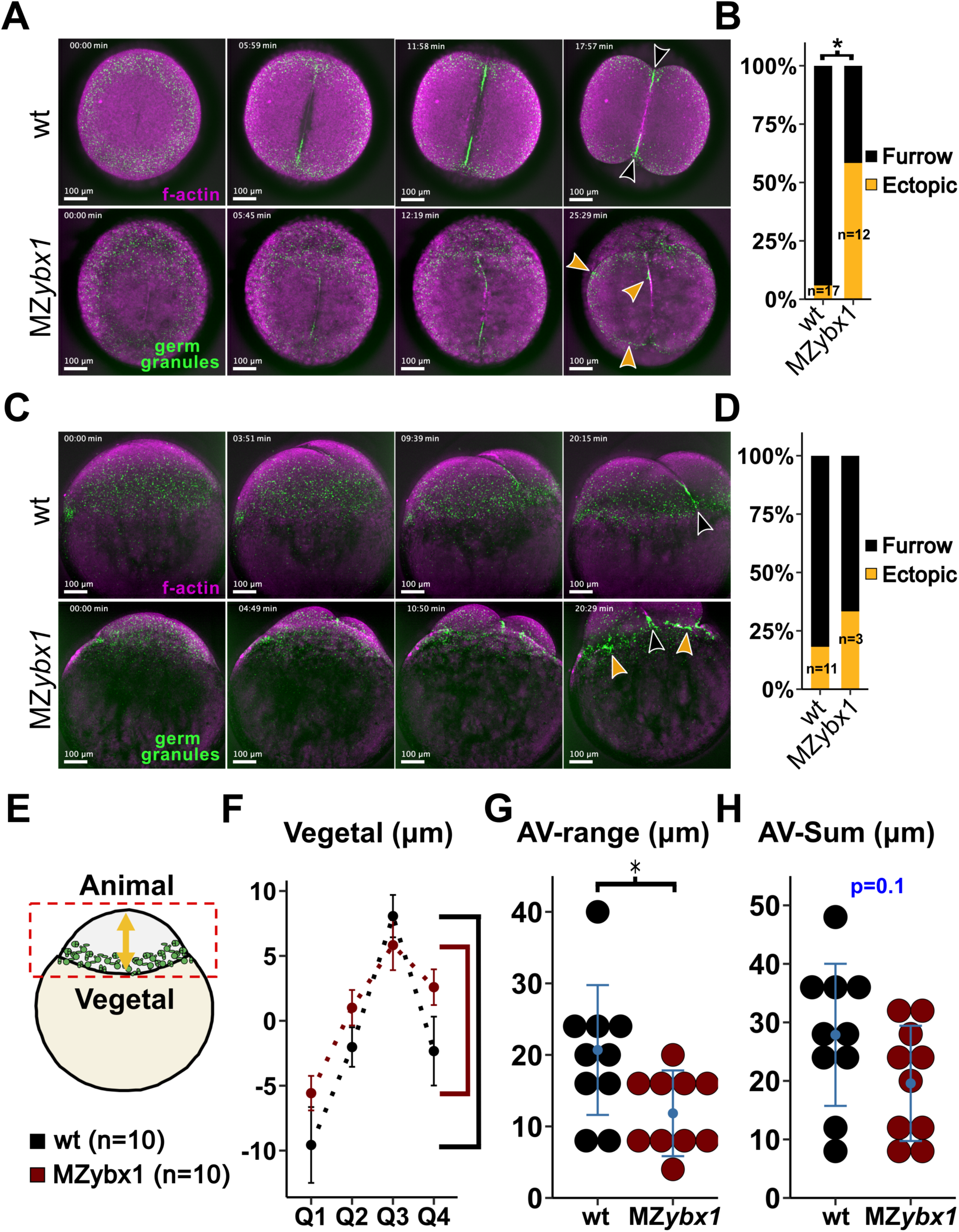
Reduced animal-ward movement of germ granules in MZ*ybx1* mutants. Stills from time-lapse imaging of Tg(buc:buc-egfp); Tg(actb2:utrn-mCherry) mutant and wt control embryos during the first cleavage division in animal pole views; Buc-EGFP reporter labels germplasm (green), Utrn-mCherry reporter labels f-actin (magenta). Black arrowheads show germplasm aggregates in the corners of the cleavage furrows in wt controls. Dark gold arrow heads indicate aberrant germplasm aggregation at the blastoderm margin and smaller furrow aggregates in the mutant. **B)** Bar chart showing % embryos with aberrant germplasm aggregation during first two cleavage divisions. **C)** Stills from time-lapse imaging of Tg(buc:buc-egfp); Tg(actb2:utrn-mCherry) mutant and wt double transgenic embryos during the first cleavage division; lateral views, animal to the top. The Buc-EGFP reporter labels germplasm in the green channel and the Utrn-mCherry reporter labels f-actin in the red channel (magenta). Black arrow heads point to furrow aggregates, and dark golden arrow heads to aberrant germplasm aggregation at the blastoderm margin **D)** Bar chart showing % embryos with aberrant germplasm aggregation during first two cleavage divisions. **E)** Schematic of 1-cell embryo in standard orientation with animal to the top. Germ granules in green, yolk is beige, and cytoplasm is grey. The extent of the embryo along the animal-vegetal axis (yellow arrow) captured in the animal-pole experiments is indicated by a dashed red box. **F)** Comparison of germ granule movement profiles in the animal-vegetal dimension in MZ*ybx1* mutants versus wt embryos. Brackets (right) indicate the range of the maximum displacement in that dimension for the two groups. **G)** Dot plot for the range of the maximum displacement in the animal-vegetal dimension in MZybx1 mutant embryos compared to wt embryos. **H)** Dot plot showing sum of displacement in the animal-vegetal dimension in MZ*ybx1* mutant embryos versus wt embryos.

Importantly, 3D tracking shows that the degree of germ granule movement along the animal-vegetal axis is reduced in MZ*ybx1* mutants compared to controls (Figure 7D and Figure 8E-H). The range of maximum movement along this axis, i.e., the range between the highest and the lowest value of the profile, is significantly lower in the furrow and corner sectors of mutants (furrow p-value = 0.02, corner p-value = 0.006). The sum of values of the animal-vegetal profile is also lower in both sector groups, but the difference is only significant for the corner sector (furrow p-value = 0.11, corner p-value = 0.05). This tendency is also present in the data for the orthogonal sectors without reaching significance (Vegetal-Range p-value = 0.18, Vegetal-Sum p-value = 0.23).

We used LLSM to complement our investigation of germ granule dynamics by acquiring movies of germ granule movements after fertilization and prior to the 1^st^ cleavage furrow formation, in wt, and MZ*ybx1* mutant embryos. (Figure S5A,B). 3D track analysis of germ granule movements (Figure S5C-I) revealed that the germ granules on average moved towards animal to a lesser extend in ybx1 mutants compared to wt controls (Figure S5D).

In summary, germ granules ectopically aggregate at the blastoderm margin with a higher frequency in *ybx1* mutants than in wt controls. Our analysis of germ granule dynamics from SDM and LLSM imaging data revealed reduced movement along the AV-axis in *ybx1* mutants. This difference in germ granule dynamics between *ybx1* mutant and wt embryos is more pronounced in the furrow region.

## Discussion

We developed a framework to study germ granule dynamics in early zebrafish embryos through quantitative approaches. This framework comprises of a set of tools for the measurement of parameters of germ granule dynamics from 3D movies acquired on a variety of microscopes. We used custom-made 3D-printed tools to mount zebrafish embryos for imaging on inverted (SDM, LLSM) and upright (wide field) microscopes. Parameter measurement from time-lapse imaging data was done using image analysis software run in MATLAB combined with custom-made scripts (Fiji, MATLAB, R) for in depth analysis. PIV lab software was used to extract the vectors of the flow of germ granules, and u-track3D was used to obtain 3D coordinates of germ granules across frames.

We used this set of tools to quantitatively characterize germ granule dynamics in normally developing embryos expressing the germplasm reporter Buc-Egfp (Figures1-4). Our work produced several insights:

1. Germ granule behavior is highly stereotypic for different populations of germ granules, based on their position relative to the cleavage furrow.
2. Germ granules move towards vegetal along cortical paths outside of the furrow, and ingress together with the furrow.
3. Bulk dynamic changes in the distribution of germ granules only commence following furrow initiation.

A proposed model for the mechanism of germ granule aggregation during the first cleavage division suggests that the embryo coopts the cell division machinery for this purpose and links germ granule dynamics to the ones of the actin and microtubular cytoskeletal networks (Eno and Pelegri, 2013). Association of germ granules with f-actin and the ends of growing microtubules (Eno and Pelegri, 2018; Theusch et al., 2006), and the interaction of germ granules with factors that bridge these cytoskeletal networks (Nair et al., 2013) provide a physical basis for the linkage. In this model most of the granule movements are driven by growing microtubules plus ends pushing on assemblies of f-actin and germ granules.

Our data is not fully compatible with the above model. We recorded only minimal changes in the distribution of germ granules in the minutes prior to the formation of the cleavage furrow (Movies2,5,7). Rather, the start of significant movement of germ granules within and outside of the furrow region coincides with the appearance of the furrow itself (Figure 4, Movies2,5,7). The first cleavage furrow forms after the 1^st^ mitotic spindle has fully formed, and before formation of the second mitotic spindle. Hence, there is a discrepancy between the timing of germ granule movement during the 1^st^ division and the timing of mitotic spindle growth. Previous work showed that zebrafish mutants such as *ack* and *cei* form the 1^st^ mitotic spindle, but fail to aggregate germplasm (Kishimoto et al., 2004; Yabe et al., 2009). Another model proposes that in early Xenopus and zebrafish embryos, where cells are unusually large, astral microtubules in metaphase are too short for spindle orientation and positioning, and that dynein pulling forces act before astral microtubules contact the cortex to orient centrosomes(Wühr et al., 2010). Consistent with this model, our observations suggest that astral microtubule growth *per se* is unlikely to be the driving force for germ granule aggregation at the distal ends of furrows.

We found that movement of furrow aggregates mimics furrow ingression during the second half of the cell division. Germ granule dynamics apparently follow dynamic changes of f-actin, which flows into the cleavage furrow during ingression. P-Myosin II was shown to be a component of germ granules, and mutant embryos for the microtubule plus end motor kinesin fail to accumulate germplasm in the corners of the cleavage furrows (Campbell et al., 2015; Nair et al., 2013). Furthermore, slow Ca^2+^ waves acting on actomyosin have been linked to germplasm dynamics (Eno et al., 2018). This could be reconciled in a model where long-range transport along microtubules brings germ granules to the regions near the cell cortex, whereafter the granules move together with cortical flow.

The current models for germplasm distribution do not consider the potential contribution of biophysical factors including cytoplasmic flow patterns or changes in the cortex in early embryos. Cytoplasmic flows can be linked to the dynamics of cytoskeletal networks via hydrostatic interactions/coupling (Pelletier et al., 2020; Shamipour et al., 2021). A similar coupling can also occur between motor-driven transport along cytoskeletal tracks and the movement of the surrounding cytoplasm (Lu and Gelfand, 2023). A recent study in C. elegans found that cortical flow coupled movement of PAR complexes into the cleavage furrow to propagate cell polarity during early embryonic cell divisions (Ng et al., 2023). Cell shape changes can also drive cytoplasmic flows (Klughammer et al., 2018), and may be active during early zebrafish development. Careful investigation of germ granule dynamics relative to other dynamic processes during early development is needed to understand what drives germ granule dynamics and distribution during early cleavage divisions.

One type of cytoplasmic flow present during early zebrafish development is ooplasmic flow that separates ooplasm from yolk after egg activation (Shamipour et al., 2019). Germ granules need to reach the blastodisc prior to recruitment to the cleavage furrow and subsequent compaction. Reduced accumulation of germplasm in the corners of the cleavage furrows together with ectopic aggregation at the blastoderm margin could be an outcome in mutants with blastodisc elevation defects. Impaired ooplasmic flow driven germ granule movement might underly the altered germplasm distribution and germline defects in *ybx1* mutants. Ooplasmic streaming is believed to be driven by actin flows. However, little is known about the link between Ybx1 and the actin cytoskeleton, and warrants further investigation.

There are certain limitations and caveats regarding our study. Most germ granules accumulate at the animal pole of mature oocytes (eggs), but germ granules also contain RNAs such as *dazl* and *bruno-like*, that localize to the vegetal cortex of eggs. It is possible that the lack of incorporation of a subset of germplasm RNPs the furrow aggregates (vegetal versus animal germplasm RNPs) may underlie the modest germline defects seen in ybx1 mutants.

Germplasm RNAs have at least two different subcellular localization patterns within early embryos: to germ granules, and to the cytoplasm outside germ granules. A shift towards cytoplasmic localization of germplasm RNAs in *ybx1* mutants would offer an alternative explanation for the reduced accumulation of germplasm RNPs in the distal cleavage furrows. Dynamic changes in the germplasm RNA solubility during early zebrafish development have recently been demonstrated (Hwang et al., 2023). Furthermore, Ybx1-binding results in translational repression of many RNAs in zebrafish embryos (Kumari et al., 2013; Sun et al., 2018; Zaucker et al., 2018; Nagorska et al., 2023). It has been proposed that poorly translated RNAs get recruited into germ granules in C elegans (Parker et al., 2020). Therefore, it is possible that changes in the subcellular localization of germplasm RNAs contribute to the germline defect in ybx1 mutant embryos.

Our framework to the study of germ granule dynamics during early zebrafish development revealed novel insights and detected subtle differences in RNP granule dynamics between *ybx1* mutants and controls. This strategy could be deployed to study the role of other processes and factors in driving germ granule movement, and for studying other RNP granules in a range of systems.

## Materials and Methods

### Zebrafish methodology

#### Husbandry and embryology

Adult zebrafish were kept at ambient temperature in the University of Warwick aquatics facility, in compliance with the University of Warwick animal welfare and ethical review board (AWERB) and the UK home office animal welfare regulations (Westerfield, 2007), covered by UK Home office licenses.

Embryos for experiments were obtained from natural spawning in 1.7 L slope tanks (Tecniplast). Egg Water (60 µg/ml “Instant Ocean” Sea Salts in dH_2_O) was used to grow embryos for raising next generations of fish, or for genotyping adult transgenics by fluorescence screening of embryos. The medium used in experiments was 0.3X Danieau’s solution (17.4 mM NaCl, 0.21 mM KCl, 0.12 mM MgSO_4_, 0.18 mM Ca(NO_3_)_2_, 1.5 mM HEPES, pH 7.6). All media were prewarmed to 28.5°C, and embryos were kept in 28.5°C incubators in between procedures.

For most experiments the “egg shell”, chorion, of the embryos was removed before the 2-cell stage, either enzymatically with Pronase or manually using fine tweezers (Dumont #5). Enzymatic dechorionation was achieved by incubation in 2.5 ml of 2 mg/ml Pronase in Egg Water for 1-3 mins. The Pronase was removed by four consecutive washes with 250 ml Egg Water in a 250 ml glass beaker. We used glass Pasteur pipettes for transferring dechorionated embryos by pipetting. Dechorionated embryos were kept in agarose-coated dishes. “Inverted” eye lash tools were used to move and orient embryos during sorting and mounting procedures, with the pointed end of the eye lash embedded in the paraffin wax.

#### Zebrafish wt and mutant strains

The work was carried out with the *ybx1 sa42* strain of zebrafish *ybx1* mutants (Kumari, 2013). The mutation had been generated and is maintained in the TU wt background. The *ybx1* mutant line was crossed into *Tg(buc:buc-egfp)* and *Tg(actb2:mCherry-Hsa.UTRN)* transgenic reporter lines (Riemer et al., 2015). The *Tg(buc:buc-egfp)* transgenic line also carried a mutation in the *buc* gene, *buc p106*, to compensate for the third copy of the *buc* gene provided by the transgene. Principally, crossing schemes were such that they produced adult females that only could be heterozygous for the transgenes and the buc p106 mutation but covered all possible *ybx1 sa42* genotypes. One family of fish used in this study originated from a cross that could yield fish homozygous for the *Tg(buc:buc-egfp)* and *Tg(actb2:mCherry-Hsa.UTRN)* transgenes. Only fish heterozygous for the transgenes were used from that cross.

#### Genotyping of mutants and transgenics

All genotyping of mutant alleles was PCR-based. DNA extraction from fin clips was performed as described (Vong et al., 2021). The *ybx1 sa42* mutation generates an additional AluI restriction site in a multiple AluI restriction site containing 502 bp region surrounding the mutation, which can be PCR amplified using primer pair sa42-fw (TTGGGGACAGTGAAATGGTT) and sa42-rev (GAGTCAAACTAAGCTACGACTAAAAGC). The largest fragment, 348 bp, from AluI digest of the respective PCR products was used to distinguish wt from sa42 mutant alleles because it is further cut into 294 bp and 54 bp fragments due to the mutation.

Genotyping of the buc p106 allele was achieved by two separate PCRs with three primers: The 412 bp PCR product from the primer “buc CAPS rev” (GAGAGAGGGGGTTGTGTC) and “buc106 wt rev” (CAGCAGTGCATAATGGTTTAG) combination serves as an internal control for the PCR reaction and suppresses by-products. The combination of primer “buc106 wt rev” with the allele specific primers “buc106 mut fw” (CATTGAGAAGGTCGACTTGA) or “buc106 wt fw” (CATTGAGAAGGTCGACTTGT) detects either the wt or the mutant allele, in separate reactions.

### Imaging

#### Imaging of germ granules by spinning disk microscopy (SDM)

Mounting of the samples for imaging on inverted microscopes was achieved as described in (Zaucker et al., 2021). The 3D-printed tools used in this study were slightly modified versions of the original tools. Time-lapse movies were acquired on a Andor Revolution XD SDM equipped with 2× Andor iXon 888 cameras; it was only used in one camera mode. The objective used was a 10× air objective (NA = 0.45, WD 4 mm). The acquisition specifics for each movie are listed in Table 1.

#### Imaging of germ granules by widefield fluorescence microscopy

Mounting of the samples for imaging on upright microscopes was achieved as described in (Zaucker et al., 2021). Z-stacks were acquired on a Nikon ECLIPSE Ni microscope equipped with a HAMAMATSU ORCA-Flash4.OLT digital camera. The objectives used were a 20× (NA = 0.5, WD 2 mm) or a 40× (NA = 0.8, WD 3.5 mm) water dipping objective.

#### Imaging of germ granules dynamics by lattice lightsheet microscopy

Samples were mounted for imaging as described in (Zaucker et al., 2021). Briefly, a single agarose well was created within the 3i LLSM sample holder by using a 3D-printed mould. We imaged germ granule movements in a small area at the blastoderm margin of wt and MZ*ybx1 sa42* mutant embryos.

Imaging was carried out on a 3i lattice lightsheet microscope (version 2) with a Coherent Sapphire 300mW 488nm laser. The annual mask had a 0.493 inner and 0.55 outer numerical aperture. The objectives were a Special Optics 0.7 NA LWD WI for excitation and a Nikon CFI Apo LWD 25× 1.1 NA WI, with a 2.5× tube lens, for detection. The emission light was filtered with a Semrock 446/523/600/677 nm quad-band filter and detected using a Hamamatsu ORCA-Flash4.0 V3 camera.

Acquisition was carried out in the Slidebook 6 software using an individual frame size of 1024×1024 pixels in X and Y (104 nm pixel size) with a 5 ms exposure time. 3D volumes consisted of 101 planes with a 1 µm step size (0.524 µm after deskewing) were acquired continuously, without interval, every 1.011 s for 300 volumes in total (∼5 mins).

### Image analysis

#### Standardized segmentation of embryos imaged in animal pole views

This was carried out in Fiji software (Schindelin et al., 2012). The imaged embryos were reoriented with the (future) cleavage furrow in horizontal orientation by rotation of the 4D images. The reoriented movies were then cropped to a size 1142×1142 px, centered on the middle of the embryo. The script “Draw Concentric Quadrants” (Olivier Burri, 2016) was used to generate a set of ROIs for concentric sectors around the center of the embryo (image):

1. A horizontal line was drawn across the center of the image (width = 1142 px, height = 1 px, y= 571 px).
2. The script output was set to 1 circle with eight quadrants
3. The set of ROIs for the sectors (quadrants) was saved

This set of ROIs was used in conjunction with a custom-made script that crops individual sectors and reorients them into the same orientation. After standardized segmentation the imaged embryo has been partitioned into eight equal-seized sectors in an embryo margin to center orientation, i.e., the wide part of the sector is at the top (circumference), and the tip of the sector (center of embryo) is at the bottom.

#### PIV analysis

The MATLAB based application PIVLab (Thielicke and Sonntag, 2021) was used for PIV analysis of the directionality of flow of germ granules. The germ granules were segmented out prior to PIV analysis by threshold segmentation of the MIPs of the movies in Fiji, to ensure that the PIVlab output is solely based on their dynamics. Slightly different thresholds for pixel values were used for the different movies, since using the same threshold was not appropriate for all movies to segment out the granules. The filtered movies were uploaded into PIVlab to generate vector maps for the flow of germ granules during the first cleavage division, and to extract vector angles from the analysis of individual sectors.

The default PIV settings were used with following Pass setup (interrogation area/Step): Pass 1 (256/128), Pass 2 (128/64), Pass 3 (64/32), Pass 4 (32/16). Velocity based validation was applied with the default settings, but without interpolation of missing data. Vector maps for the averaged movement of germ granules during the 1^st^ division were plotted by calculating the mean for all frames in the “Derive Temporal Parameters” menu. Vector angles were obtained by saving the output of plotting “Vector direction” in the “Derive Parameters” menu.

### Track analysis of the movement of germ granules in 3D

#### LLSM data

The data were processed firstly by deskewing in Slidebook 6 (imaged volume was 106 x 106 x 53 μm). Subsequently, the data were down sampled laterally (X and Y) by two-fold in Fiji and exported as 16-bit tiffs.

Tracking of the germ granules was carried out with the u-track3D software using the GUI. The default spot detection algorithm was used (called “multiscaleDetectionDebug”) with scales set to 4 and an alpha sensitivity of 0.0001. The tracking used a “maximum gap to close” of 1 frame, a “minimum length of track segments from first step” of 3 frames, with segment/track merging allowed, and segment/track splitting disallowed. The cost function for frame-to-frame linking used the “Brownian and directed motion models” with “directed motion position propagation” allowed, “instantaneous direction reversal” disallowed, a lower bound search radius of 2 pixels, an upper bound of 10 pixels, and used 2 frames for nearest neighbour distance to expand the Brownian search radius. The cost function gap closing, merging and splitting used the “Brownian and directed motion models” with the following settings: lower bound of 2 pixels, upper bound of 10 pixels, 2 frames for nearest neighbour distance to expand the Brownian search radius, 0.5 scaling power in fast expansion phase to expand the search radius with gap length, 0.01 scaling power in slow expansion phase, gap length of 1 to transition from fast to slow expansion, penalty of 1.5 for increasing gap length, the allowed ratio of intensities before and after merging were a minimum of 0.5 and a maximum of 2, for motion modelling the minimum track segment lifetime for classification as linear or random was set to 5 frames, the multiplication factor for linear search radius was 1, and the linear motion search radius was scaled with time using a scaling factor of 1 in the fast expansion phase, and 0.01 in the slow expansion phase, with a gap length of 1 to transition from the fast to slow expansion phase, the maximum angle between linear track segments was 30 degrees, the birth and death cost was set to automatically determined using percentiles. The Kalman filter functions were set to Brownian and directed motion models using the default settings. The detection and tracking settings were optimised by visual comparison to manually detected and tracked data.

#### SDM data

Germ granules were tracked with u-track3D software (Roudot et al., 2023) run in MATLAB. For each movie the individual sectors from standardized segmentation were imported into u-track3D software together with the corresponding metadata via the software’s GUI. The tracking software was then run on the full list of eight sectors. The settings differed from the ones used for the LLSM data:

Spot detection scales were set to 3 - 5 - 7 and an alpha sensitivity to 0.05. In the settings for frame-to-frame linking “instantaneous direction reversal” was allowed, the lower bound search radius was 1 pixel, the upper bound 7 pixels. In gap closing, merging and splitting settings lower bound was 1 pixel, the upper bound 7 pixels, for motion modelling the multiplication factor for linear search radius was 3, and the linear motion search radius was scaled with time using a scaling factor of 0.5 in the fast expansion phase.

### Parameters of germ granule dynamics from track analysis

#### LLSM data

The u-track3D tracking data were reformatted into a table (csv file) using a custom-made MATLAB script. A custom-made R script was used to calculate parameters of germ granule movement from the tracking data (3D coordinates) on the table. The script can return parameters for any time interval within the 5 min movies, which allows for timewise alignment of movie segments from different embryos. We used this feature to compare parameters of germ granule movements in 1 min intervals, from 21 minutes post fertilization (mpf) onwards. The 1st cleavage furrow, which starts to form around 35 mpf, has a dramatic impact on the movement of germ granules. To avoid the confounding impact of differences in the distance of the imaged area to forming cleavage furrows we excluded movies/segments beyond 32 mpf from the analysis. Only complete tracks for 1 min movie segments were analyzed.

We imaged two sets of *ybx1* mutant embryos and controls: Tg(buc:buc-egfp); *ybx1^sa42/sa42^*; *buc^p106/+^* double mutant transgenics, and single mutant transgenics without endogenous *buc* mutations. Only, the single mutant transgenic data was analyzed because the wt control embryos for the double mutant transgenics were all obtained from the same female, and the double mutant transgenic data did not cover the whole time period for all genotypes.

#### SDM data

A custom-made script was used to calculate parameters of germ granule dynamics for four consecutive segments, named Q1-4, of the time interval spanning the 1^st^ cleavage division. The number of frames per segment was either exactly the same, or differed by maximum one frame between different segments, dependent on whether the number of frames during the 1^st^ division was a multiple of four.

### Format of output tables with parameters

The two different R-scripts produced three output tables for the calculated parameters per time interval, i.e. 1 min interval for the LLSM data, and one of the four segments for the SDM data. To facilitate exact calculations of the speed of movement, exact frame rates for consecutive frames were used based on the corresponding timestamps in the metadata. The output tables had following format:

1. Parameters (average for each track) track_index, n (frames), Length (μm), Displacement (μm), Straightness, Speed (μm/s), Angle (degrees), Lateral {displacement in x (μm)}, Center {displacement in y (μm)}, Vegetal {displacement in z (μm)}
2. Average (average for the interval, i.e., for all tracks in the parameters table)
3. Steps (parameters for each step of all tracks of the interval)

### Vegetal-Range and Vegetal-Sum

We calculated these parameters from the profiles for the displacement along the animal-vegetal axis for individual sectors:

- Vegetal-Range = Maximum value (any segment) – Minimum value (any segment)
- Vegetal-Sum = Sum of absolute values from all four segments

## Line Scans

Line Scans were carried out in Fiji software. The imaged embryos were reoriented into the same orientation with a vertically oriented 1^st^ cleavage furrow. A 100 μm wide horizontal line that runs through the germ granule free centre of the embryo and spans the whole animal pole was generated by rectangular box selection. The data of the plot profile (line scan) of gray values was saved for further processing. A custom-made R script was used to normalize the line scan data into 0.1% bins of the total line length, which varied between embryos. Line graphs for the average pixel values across all embryos of a group (controls vs mutants) were generated in R.

## Germplasm volume measurements

Dechorionated Tg(buc:buc-egfp) transgenic embryos were mounted 6×6 arrays for imaging on an upright microscope as described (Zaucker et al., 2021). At the 2-cell stage z-stacks of the animal pole of the embryos were acquired on a Nikon widefield microscope at 20×. The image processing/analysis was carried out using custom script in Fiji. We deconvolved the data using the “Tikhonov Regularization Inverse Filter” algorithm (λ = 2.154E-04) of the DeconvolutionLab2 plugin with a virtual PSF that was generated by the PSF generator (Born & Wolf 3D Optical Model). The 3D Objects Counter plugin was used with the deconvolved data to obtain the volumes of continuous objects after threshold segmentation.

## PGCs in somitogenesis stage embryos

MZybx1 embryos were obtained from crosses of Tg(buc:buc-egfp); ybx1^sa42/sa42^ females with ybx1^sa42/sa42^ males. Control embryos were obtained from crosses of Tg(buc:buc-egfp); ybx1^sa42/+^ females with wt males. All embryos were shifted to 22℃ at 30% epiboly and incubated until imaging at somitogenesis. The gonadal region of different groups of embryos was imaged in dorsal views on a Nikon widefield microscope during three consecutive periods: 3 to 6-somite, 6 to 10-somite, 10 to 14-somite. The number of PGCs per embryo was counted visually by scanning through the acquired z-stacks. Only the 10 to 14-somite data was analyzed because it is close to the stage when all PGCs have arrived in the gonadal region (15-somite).

## Comparative Gene Expression Analysis

### Whole mount in situ hybridization (WISH)

The ISH experiments against *ddx4* and *dnd1* were carried out using standard protocols as described in (Lim et al., 2012).

### qRT-PCR

375µl of RNA protectant was added once to approximately 25 embryos and then homogenised with a needle and syringe until there were no pieces left and then flash frozen in liquid nitrogen and stored in -80°C. Proteinase K reaction buffer (30µl) was added and Proteinase K (15µl) at 55°C for 5 minutes. The mixture was mixed and spun down for 2 minutes at 16000×g. The supernatant was then transferred, and an equal volume of lysis buffer was added and vortexed. For RNA purification, 800µl of the sample from the extraction was placed in a gDNA column fitted with a collection tube and spun for 30 seconds. The gDNA column was then discarded. An equal volume of 100% ethanol was added to the flow-through and mixed thoroughly by pipetting. The mixture was then added to an RNA Purification column and fitted with a collection tube and spun for 30seconds and the flow-through was discarded. 500µl of RNA wash buffer was added and spun for 30seconds and the flow-through was discarded. In a RNase-free tube, 5µl of DNase I and DNase I Reaction Buffer was pipetted directly on top of the column matrix. This mixture was then incubated for 30-45minutes. 500µl of RNA Priming Buffer was added and spun for 30seconds, and the flow-through was discarded. RNA Wash Buffer (500µl) was added and spun for 30seconds and the flow-through was again discarded. Another 500µl RNA wash buffer was added and spun for 2 minutes, and the column was then transferred to the RNase-free tube and re-spun for 1minute. 30-100µl Nuclease-free water was added to the centre of the column matrix and spun for 30 seconds. Next 1µl of glycogen and 15µl of ammonium acetate stop solution was added, followed by ethanol precipitation and storage at -80°C. Following measurement of concentration by Nanodrop and gel electrophoresis to assess quality, first strand cDNA synthesis was performed according to the Thermo Fisher™ Protocol. PCR was performed as per the2× qPCRBIO SyGreen Blue Mix Lo-ROX™ Protocol.

### RNA-Seq

Samples of P*ybx1* and M*ybx1* embryos at the 4-cell stage were lysed, total RNA was purified. Following quality and concentration analysis by gel electrophoresis, nanodrop and Bioanalyzer measurements, libraries were prepared for next-gen sequencing (NGS). Standard controls were included to ensure high quality library preparation. Low quality reads were filtered out and reads that aligned to the zebrafish reference genome assembly and Ensembl protein coding gene models were analyzed. The CDS reads/genic reads ratio and RPKM counts were compared between P*ybx1* and M*ybx1* mutants.

### Statistical Tests

The statistic software R-Studio was used for descriptive (graphs) and inferential statistics (statistical hypothesis testing). Parametric tests were used for normally distributed data, and non-parametric tests for any other data. Shapiro-Wilk tests were performed to test for normal distribution. For testing differences in the means of one dependent variable between two groups (independent variable), t-tests or Mann-Whitney U tests were used at a significance level of 5% (p-value of 0.05). For testing differences in the means of one variable dependent on the values of two independent variables (e.g., time and group), two-way ANOVA in conjunction with Tukey’s post hoc tests at a significance level of 5% (p-value 0.05) were used. Fisher Exact tests at a significance level of 5% (p-value 0.05) were used to test for significant relationships between categorical variables (contingency tables).

## Supporting information

Zaucker supplemental legends

Zaucker supplemental figures

## Acknowledgements

We thank Roland Dosch for the Tg(buc:buc-egfp) line; members of the Sampath laboratory and members of the Warwick fish community for feedback and suggestions; MP is supported by a Warwick ARAP doctoral scholarship, DSLR was supported by the BBSRC MIBTP, BL was supported by the MRC DTP; Work in the Sampath laboratory is supported by funds from the Leverhulme trust and the BBSRC. The lattice light sheet microscope was funded by a Wellcome Trust Multi-User Equipment Grant 208384/Z/17/Z. We thank Laura Cooper of CAMDU (Computing and Advanced Microscopy Unit) and Lewis Mosby for their support & assistance with MATLAB scripts.

## References

Boke, E., Ruer, M., Wühr, M., Coughlin, M., Lemaitre, R., Gygi, S. P., Alberti, S., Drechsel, D., Hyman, A. A. and Mitchison, T. J. (2016). Amyloid-like Self-Assembly of a Cellular Compartment. Cell 166, 637–650.

Campbell, P. D., Heim, A. E., Smith, M. Z. and Marlow, F. L. (2015). Kinesin-1 interacts with bucky ball to form germ cells and is required to pattern the zebrafish body axis. Development (Cambridge) 142,.

Chen, R., Stainier, W., Dufourt, J., Lagha, M. and Lehmann, R. (2024). Direct observation of translational activation by a ribonucleoprotein granule. Nat Cell Biol 26, 1322–1335.

Chernov, K. G., Mechulam, A., Popova, N. V., Pastre, D., Nadezhdina, E. S., Skabkina, O. V., Shanina, N. A., Vasiliev, V. D., Tarrade, A., Melki, J., et al. (2008). YB-1 promotes microtubule assembly in vitro through interaction with tubulin and microtubules. BMC Biochem 9,.

Czolowska, R. (1969). Observations on the origin of the “germinal cytoplasm” in Xenopus laevis. J Embryol Exp Morphol 22,.

Deis, R., Kar, S., Ahmad, A., Bogoch, Y., Dominitz, A., Shvaizer, G., Sasson, E., Mytlis, A., Ben-Zvi, A. and Elkouby, Y. M. (2022). Multi-Step Cellular Control of Molecular Condensation by Microtubules in Early Oogenesis. BioRχiv preprint,.

Elkouby, Y. M., Jamieson-Lucy, A. and Mullins, M. C. (2016). Oocyte Polarization Is Coupled to the Chromosomal Bouquet, a Conserved Polarized Nuclear Configuration in Meiosis. PLoS Biol 14,.

Eno, C. and Pelegri, F. (2013). Gradual recruitment and selective clearing generate germ plasm aggregates in the zebrafish embryo. Bioarchitecture 3,.

Eno, C. and Pelegri, F. (2018). Modulation of F-actin dynamics by maternal Mid1ip1L controls germ plasm aggregation and furrow recruitment in the zebrafish embryo. Development (Cambridge) 145,.

Eno, C., Gomez, T., Slusarski, D. C. and Pelegri, F. (2018). Slow calcium waves mediate furrow microtubule reorganization and germ plasm compaction in the early zebrafish embryo. Development (Cambridge) 145,.

Hashimoto, Y., Maegawa, S., Nagai, T., Yamaha, E., Suzuki, H., Yasuda, K. and Inoue, K. (2004). Localized maternal factors are required for zebrafish germ cell formation. Dev Biol 268,.

Hwang, H., Chen, S., Ma, M., Divyanshi, Fan H. C., Borwick, E., Böke, E., Mei, W. and Yang, J. (2023). Solubility phase transition of maternal RNAs during vertebrate oocyte-to-embryo transition. Dev Cell 58,.

Illmensee, K. and Mahowald, A. P. (1974). Transplantation of posterior polar plasm in Drosophila. Induction of germ cells at the anterior pole of the egg. Proc Natl Acad Sci U S A 71,.

Kawaguchi, A., Asaka, M. N., Matsumoto, K. and Nagata, K. (2015). Centrosome maturation requires YB-1 to regulate dynamic instability of microtubules for nucleus reassembly. Sci Rep 5,.

Kishimoto, Y., Koshida, S., Furutani-Seiki, M. and Kondoh, H. (2004). Zebrafish maternal-effect mutations causing cytokinesis defect without affecting mitosis or equatorial vasa deposition. Mech Dev 121,.

Klughammer, N., Bischof, J., Schnellbächer, N. D., Callegari, A., Lénárt, P. and Schwarz, U. S. (2018). Cytoplasmic flows in starfish oocytes are fully determined by cortical contractions. PLoS Comput Biol 14,.

Knaut, H., Pelegri, F., Bohmann, K., Schwarz, H. and Nüsslein-Volhard, C. (2000). Zebrafish vasa RNA but Not Its Protein Is a Component of the Germ Plasm and Segregates Asymmetrically before Germline Specification. J Cell Biol 149, 875–888.

Kumari, P., Gilligan, P. C., Lim, S., Tran, L. D., Winkler, S., Philp, R. and Sampath, K. (2013). An essential role for maternal control of Nodal signaling. Elife 2, e00683.

Lim, S., Kumari, P., Gilligan, P., Quach, H. N. B., Mathavan, S. and Sampath, K. (2012). Dorsal activity of maternal squint is mediated by a non-coding function of the RNA. Development 139, 2903–2915.

Lu, W. and Gelfand, V. I. (2023). Go with the flow – bulk transport by molecular motors. J Cell Sci 136,.

Mehta, S., Algie, M., Al-Jabry, T., McKinney, C., Kannan, S., Verma, C. S., Ma, W., Zhang, J., Bartolec, T. K., Masamsetti, V. P., et al. (2020). Critical role for cold shock protein YB-1 in cytokinesis. Cancers (Basel) 12,.

Moravec, C. E. and Pelegri, F. (2020). The role of the cytoskeleton in germ plasm aggregation and compaction in the zebrafish embryo. In Current Topics in Developmental Biology,.

Nair, S., Marlow, F., Abrams, E., Kapp, L., Mullins, M. C. and Pelegri, F. (2013). The Chromosomal Passenger Protein Birc5b Organizes Microfilaments and Germ Plasm in the Zebrafish Embryo. PLoS Genet 9,.

Ng, K. B., Hirani, N., Bland, T., Borrego-Pinto, J., Wagner, S., Kreysing, M. and Goehring, N. W. (2023). Cleavage furrow-directed cortical flows bias PAR polarization pathways to link cell polarity to cell division. Current Biology 33,.

Parker, D. M., Winkenbach, L. P., Boyson, S., Saxton, M. N., Daidone, C., Al-Mazaydeh, Z. A., Nishimura, M. T., Mueller, F. and Nishimura, E. O. (2020). mRNA localization is linked to translation regulation in the Caenorhabditis elegans germ lineage. Development (Cambridge) 147,.

Pelletier, J. F., Field, C. M., Fürthauer, S., Sonnett, M. and Mitchison, T. J. (2020). Co-movement of astral microtubules, organelles and f-actin by dynein and actomyosin forces in frog egg cytoplasm. Elife 9,.

Riemer, S., Bontems, F., Krishnakumar, P., Gömann, J. and Dosch, R. (2015). A functional Bucky ball-GFP transgene visualizes germ plasm in living zebrafish. Gene Expression Patterns 18,.

Ripin, N. and Parker, R. (2023). Formation, function, and pathology of RNP granules. Cell 186,.

Roudot, P., Legant, W. R., Zou, Q., Dean, K. M., Isogai, T., Welf, E. S., David, A. F., Gerlich, D. W., Fiolka, R., Betzig, E., et al. (2023). u-track3D: Measuring, navigating, and validating dense particle trajectories in three dimensions. Cell Reports Methods 3,.

Ruzanov, P. V., Evdokimova, V. M., Korneeva, N. L., Hershey, J. W. B. and Ovchinnikov, L. P. (1999). Interaction of the universal mRNA-binding protein, p50, with actin: A possible link between mRNA and microfilaments. J Cell Sci 112,.

Sato, K., Sakai, M., Ishii, A., Maehata, K., Takada, Y., Yasuda, K. and Kotani, T. (2022). Identification of embryonic RNA granules that act as sites of mRNA translation after changing their physical properties. iScience 25,.

Schindelin, J., Arganda-Carreras, I., Frise, E., Kaynig, V., Longair, M., Pietzsch, T., Preibisch, S., Rueden, C., Saalfeld, S., Schmid, B., et al. (2012). Fiji: An open-source platform for biological-image analysis. Nat Methods 9,.

Shamipour, S., Kardos, R., Xue, S. L., Hof, B., Hannezo, E. and Heisenberg, C. P. (2019). Bulk Actin Dynamics Drive Phase Segregation in Zebrafish Oocytes. Cell 177,.

Shamipour, S., Caballero-Mancebo, S. and Heisenberg, C. P. (2021). Cytoplasm’s Got Moves. Dev Cell 56,.

Strome, S. and Wood, W. B. (1982). Immunofluorescence visualization of germ-line-specific cytoplasmic granules in embryos, larvae, and adults of Caenorhabditis elegans. Proc Natl Acad Sci U S A 79,.

Sun, J., Yan, L., Shen, W. and Meng, A. (2018). Maternal Ybx1 safeguards zebrafish oocyte maturation and maternal-to-zygotic transition by repressing global translation.

Theusch, E. V., Brown, K. J. and Pelegri, F. (2006). Separate pathways of RNA recruitment lead to the compartmentalization of the zebrafish germ plasm. Dev Biol 292,.

Thielicke, W. and Sonntag, R. (2021). Particle Image Velocimetry for MATLAB: Accuracy and enhanced algorithms in PIVlab. J Open Res Softw 9,.

Trcek, T., Grosch, M., York, A., Shroff, H., Lionnet, T. and Lehmann, R. (2015). Drosophila germ granules are structured and contain homotypic mRNA clusters. Nat Commun 6,.

Trcek, T., Douglas, T. E., Grosch, M., Yin, Y., Eagle, W. V. I., Gavis, E. R., Shroff, H., Rothenberg, E. and Lehmann, R. (2020). Sequence-Independent Self-Assembly of Germ Granule mRNAs into Homotypic Clusters. Mol Cell 78,.

Tzung, K. W., Goto, R., Saju, J. M., Sreenivasan, R., Saito, T., Arai, K., Yamaha, E., Hossain, M. S., Calvert, M. E. K. and Orbán, L. (2015). Early depletion of primordial germ cells in zebrafish promotes testis formation. Stem Cell Reports 4,.

Uchiumi, T., Fotovati, A., Sasaguri, T., Shibahara, K., Shimada, T., Fukuda, T., Nakamura, T., Izumi, H., Tsuzuki, T., Kuwano, M., et al. (2006). YB-1 is important for an early stage embryonic development: Neural tube formation and cell proliferation. Journal of Biological Chemistry 281,.

Vong, Y. H., Sivashanmugam, L., Leech, R., Zaucker, A., Jones, A. and Sampath, K. (2021). The RNA-binding protein Igf2bp3 is critical for embryonic and germline development in zebrafish. PLoS Genet 17,.

Wang, J. T., Smith, J., Chen, B. C., Schmidt, H., Rasoloson, D., Paix, A., Lambrus, B. G., Calidas, D., Betzig, E. and Seydoux, G. (2014). Regulation of RNA granule dynamics by phosphorylation of serine-rich, intrinsically disordered proteins in C. elegans. Elife 3,.

Westerich, K. J., Tarbashevich, K., Schick, J., Gupta, A., Zhu, M., Hull, K., Romo, D., Zeuschner, D., Goudarzi, M., Gross-Thebing, T., et al. (2023). Spatial organization and function of RNA molecules within phase-separated condensates in zebrafish are controlled by Dnd1. Dev Cell 58,.

Wu, Y., Han, B., Gauvin, T. J., Smith, J., Singh, A. and Griffin, E. E. (2019). Single-molecule dynamics of the P granule scaffold MEG-3 in the Caenorhabditis elegans zygote. Mol Biol Cell 30,.

Wühr, M., Tan, E. S., Parker, S. K., Detrich, H. W. and Mitchison, T. J. (2010). A model for cleavage plane determination in early amphibian and fish embryos. Current Biology 20,.

Yabe, T., Ge, X., Lindeman, R., Nair, S., Runke, G., Mullins, M. C. and Pelegri, F. (2009). The maternal-effect gene cellular island encodes Aurora B kinase and is essential for furrow formation in the early zebrafish embryo. PLoS Genet.

Zaucker, A., Mitchell, C. A., Coker, H. L. E. and Sampath, K. (2021). Tools to Image Germplasm Dynamics During Early Zebrafish Development. Front Cell Dev Biol 9,.

