## Supplementary material for "In toto imaging of germ plasm dynamics reveals an essential role for early distribution of germ granules in germline development": Zaucker supplemental legends

**Zaucker et al.,**
**Supplemental Figure Legends**

**FigureS1 Repeat of 3d tracking experiments with single transgenics**
**A)** Quantitation of the directionality of germ granule across different sector groups based upon proportion of vectors for each category. The legend (right) explains the different categories with corresponding ranges for the vector angles and color codes: margin (slate-grey), diagonal up (sky blue), lateral (green), diagonal down (pink), center (magenta). Movement towards the margin (margin, diagonal up) was defined as "Dispersal" (light grey), whereas movement towards the center (diagonal down, center) or into the furrow (lateral) was defined as "Aggregation" (purple). The stacked bar chart on the left compares the proportion of Dispersal vs Aggregation germ granule movements between the different sector groups. Stacked bar chart (right) compares proportions of vectors for all categories of directionality across the sector groups. **B-I)** Temporal profiles for parameters of germ granule dynamics.

**Figure S2 The germline defect is a maternal effect *ybx1 sa42* mutant phenotype**
**A)** Schematic outline of temperature shift experiments with *ybx1* mutants for the temperature sensitive *sa42* allele. Temperature shifts to the restrictive temperature 22°C were performed at 30% epiboly. In some experiments, the embryos were consequently used for ISH against the germplasm marker *ddx4*, in other experiments they were grown to adulthood for the analysis of sex ratios. **B)** Sex ratios observed in fish grown from temperature-shifted embryos. **C)** Representative examples for wt and MZ*ybx1* mutant 21-somite embryos showing *ddx4* RNA (purple) expression by ISH after temperature-shift. Scale bar = 100 µm. **D)** Dot plot for the quantification of the number of PGCs per embryo for the experiment in C. **E)** Representative examples for P*ybx1* and M*ybx1* mutant 21-somite embryos stained for *ddx4* RNA (purple) by ISH after temperature-shift, compared to embryos that had not have been shifted. **F)** PGCs in the gonadal region of 10 to 14-somite stage Tg(buc:buc-egfp) embryos that contain Buc-EGFP labelled germplasm. The dashed white lines outline the midline. Scale bar = 50 µm. **G)** Dot plot comparing the number of PGCs per embryo from the experiment in E between controls (heterozygote females crossed with wt males) and MZ*ybx1* embryos.

**Figure S3 Reduced amount of Buc-EGFP labelled germplasm and PGCs in Tg(buc:buc-egfp) transgenic MZ*ybx1* mutants**
**A)** Tg(buc:buc-egfp) embryos imaged in animal pole views on a widefield microscope. Controls are embryos from $ybx^{sa42/+}$ females crossed with wt males. Scale bar = 100 µm. **B)** Dot plot comparing the volume of EGFP+ germplasm furrow aggregates per embryo for the experiment in A. **C)** Stills from time-lapse imaging of Buc-EGFP+ germplasm in control ($ybx^{sa42/+}$ mother) and MZ*ybx1* mutant embryos at the 2-cell, 4-cell and 8-cell stage. **D)** Dot plot comparing the volume of EGFP+ germplasm in the blastoderm per embryo from the experiment in C between controls and MZybx1 mutants.

**Figure S4 Speed of germ granule movement corrected for differences in the frame rate**

Line chart for the average speed profile of germ granule movement during the first cleavage divisions. A subset of the embryos from Figure 7 with highly similar frame rates was compared in this figure (average frame interval: wt = 28 s, MZ*ybx1* = 27.6 s).

**Figure S5 Movement towards animal prior to the 1st division is reduced in MZ*ybx1* mutants**

**A)** Lateral view schematics illustrating progressive distension of the blastodisc due to ooplasmic streaming during the time period that has been investigated in the experiment. **B)** Outline of LLSM experiments to measure parameters of germ granule dynamics between wt controls and MZ*ybx1* mutants before furrow formation (21-32 mpf). The movement of germ granules in a small area near blastoderm margin, indicated by a red box in the schematic of an embryo with Buc-EGFP labelled germplasm, was recorded by time lapse imaging on a LLSM. **C)** Bar plot for the average movement of germ granules in the animal-vegetal dimension at different time-points. **D)** Bar plot for the average movement of germ granules in the animal-vegetal dimension across all time-points. **E-I)** Bar plot for the comparison of other parameters of germ granule kinetics between wt siblings (black) and MZ*ybx1* (gold) across all time-points.
