## Supplementary material for "In toto imaging of germ plasm dynamics reveals an essential role for early distribution of germ granules in germline development": Zaucker supplemental figures

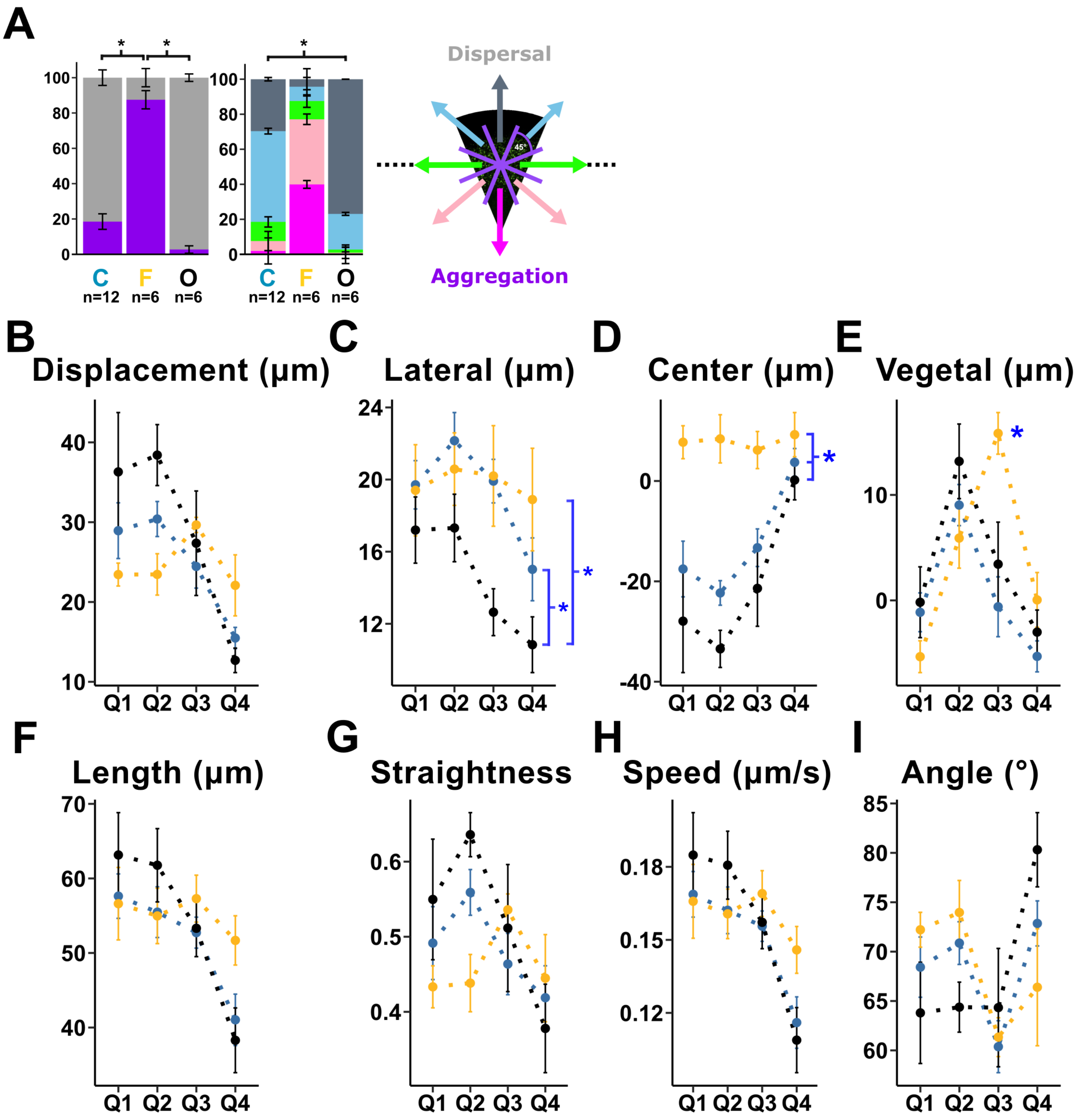


Zaucker *et al.* Figure S1

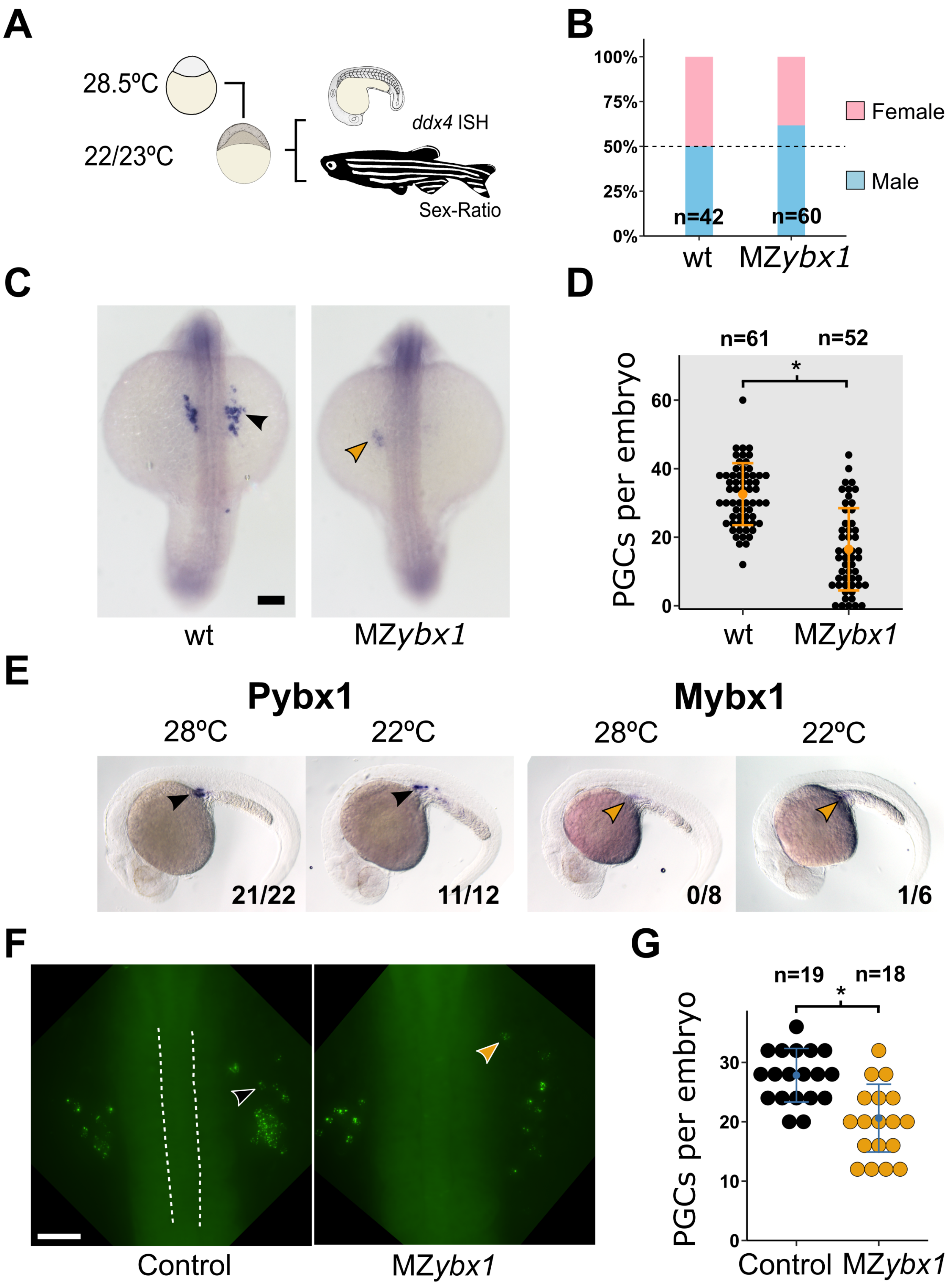


Zaucker *et al.* Figure S2

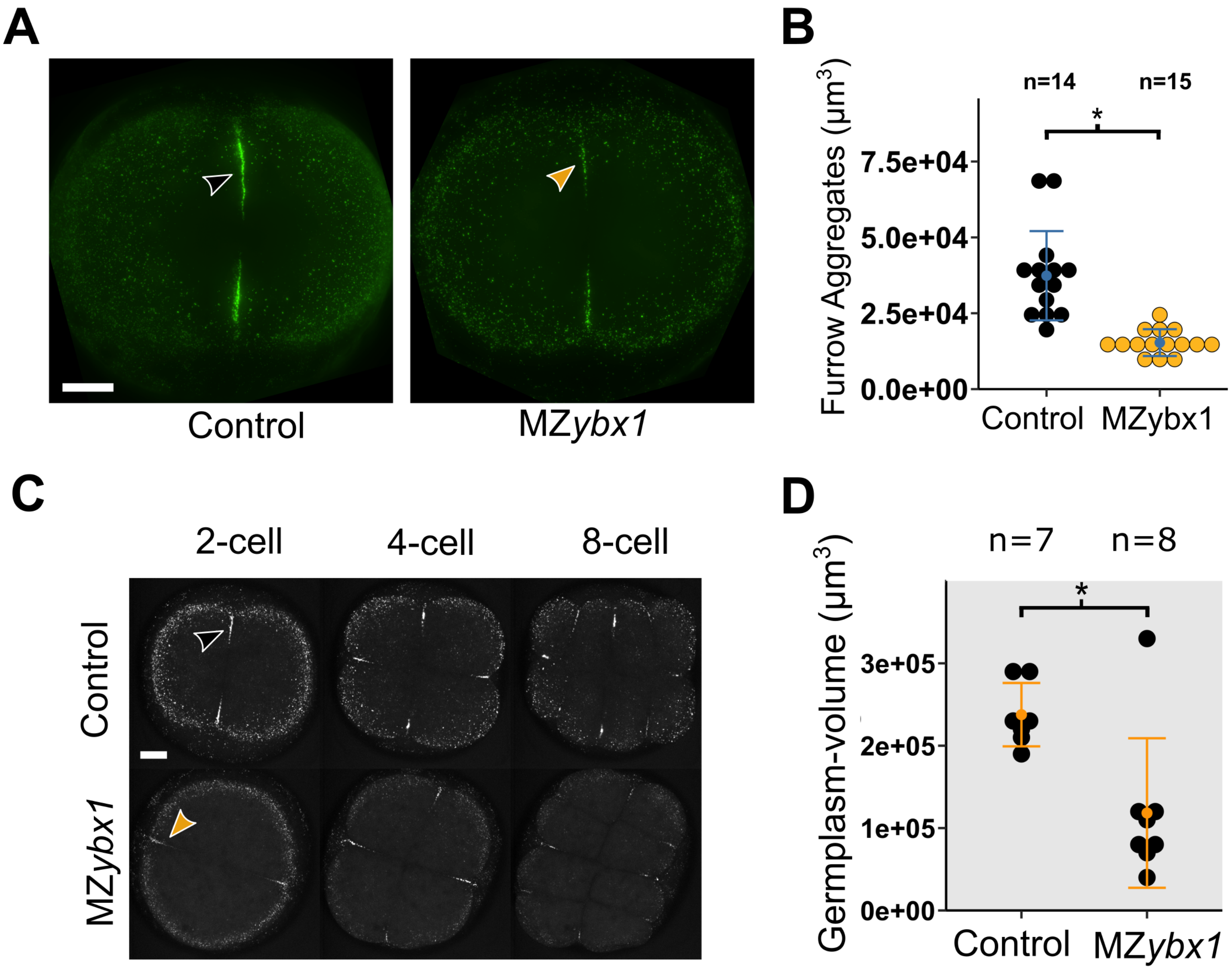


Zaucker *et al.* Figure S3

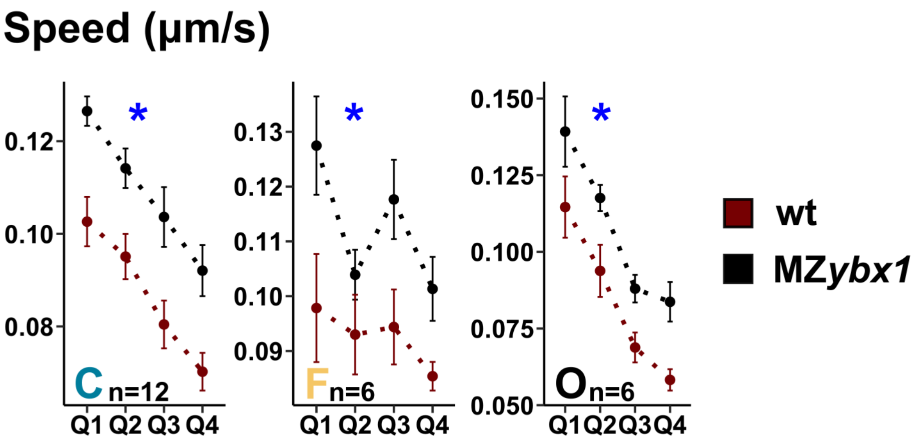


Zaucker *et al.* Figure S4

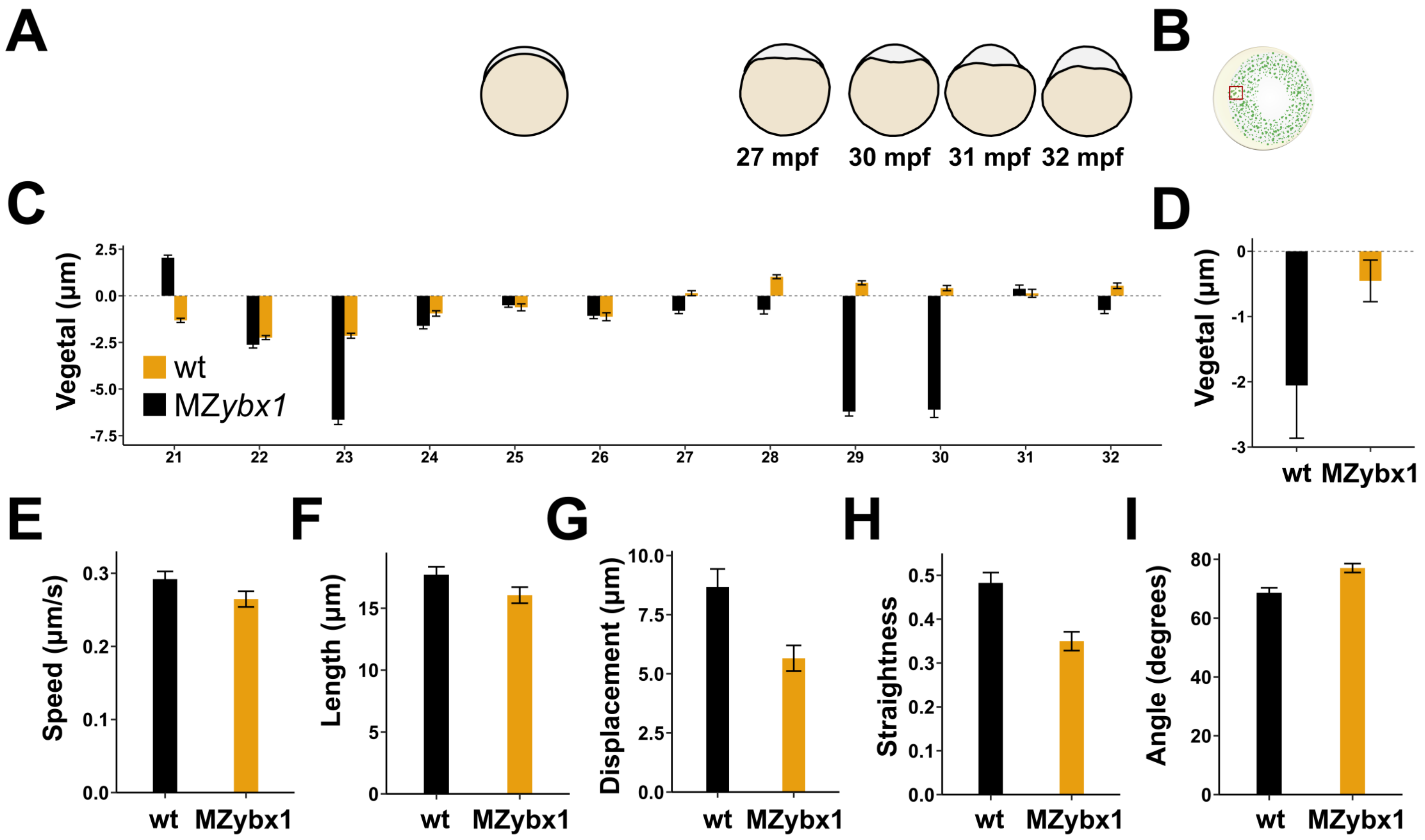


Zaucker *et al.* Figure S5
